# Molecular evolution of the meiotic recombination pathway in vertebrates

**DOI:** 10.1101/2024.06.28.599605

**Authors:** Taylor Szasz-Green, Katherynne Shores, Vineel Vanga, Luke Zacharias, Andrew K. Lawton, Amy L. Dapper

## Abstract

Meiotic recombination is an integral cellular process, required for the production of viable gametes, and the rate at which it occurs is a fundamental genomic parameter, modulating how the genome responds to selection. Our increasingly detailed understanding of its molecular underpinnings raises the prospect that we can gain insight into trait divergence by examining the molecular evolution of recombination genes from a pathway perspective, as in mammals, where protein-coding changes in the later stages of the recombination pathway are connected to divergence in intra-clade recombination rate. Here, we leveraged increasing availability of avian and teleost genomes to reconstruct the evolution of the recombination pathway across two additional vertebrate clades: birds, which have higher and more variable rates of recombination and similar divergence times to mammals; and teleost fish, which have much deeper divergence times. We found that the rates of molecular evolution of recombination genes are highly correlated between vertebrate clades, suggesting that they experience similar selective pressures. However, recombination genes in birds were significantly more likely to exhibit signatures of positive selection, unrestricted to later stages of the pathway. There is a significant correlation between genes linked to recombination rate variation in mammalian populations and those with signatures of positive selection across the avian phylogeny, suggesting a link between selection and recombination rate. In contrast, the teleost fish recombination pathway is more highly conserved with significantly less evidence of positive selection. This is surprising given the high variability of recombination rates in this clade.

## Introduction

Although meiotic recombination, the exchange of genetic material between homologous chromosomes, is ubiquitous in sexually reproducing organisms, the rate at which it occurs varies significantly within and between species (Stapley et al. 2017b). This variation has wide-ranging consequences. Meiotic recombination is a fundamental genomic and evolutionary parameter. Within populations, recombination generates new combinations of alleles, shaping patterns of genetic variation within and between genomes and modulating the efficacy of selection (Stapley et al. 2017a). Recombination rate can also directly impact zygote viability. In most species, a minimum of one crossover event per chromosome pair is required to ensure the resulting gametes each receive the proper genetic complement (Hassold and Hunt 2001).

Proximally, recombination rate is determined by the number of crossover events, which arise through a highly-regulated and complex cellular pathway. While there are still large gaps in our understanding of how and why recombination diverges between species, it is likely that variation in the genes that regulate meiotic recombination pathway are important contributors. Thus, one strategy for understanding how species diverge is to inspect molecular evolution of genes involved in this pathway.

The meiotic recombination pathway can be divided up into discrete steps. Previous research in mice, yeast, and nematodes identified roughly six stages in the meiotic recombination pathway and key genes acting on each step (Figure 1). In mammals, two lines of evidence connect protein-coding changes in later stages of the pathway to differences in rates of recombination. First, within populations, individual variation in recombination rate is linked to genetic variation at key loci in the recombination pathway (Dumont et al. 2009). Genetic variants that impact recombination rate within species have been identified in cattle, sheep, and red deer (Johnston et al. 2016; Kadri et al. 2016; Petit et al. 2017; Johnston et al. 2018; Sandor et al. 2012; Kong et al. 2008). The majority of these gene variants are found in the latter half of the recombination pathway, regulating which double strand breaks are resolved as crossovers. Second, Dapper and Payseur (2019) identified signatures of positive selection in a focal panel of meiotic recombination genes within mammals (Dapper and Payseur 2019). Specifically, genes involved with the stabilization of homologous pairing and the crossover/non-crossover (CO/NCO) decision showed greater likelihood of signatures of adaptive evolution, supporting the hypothesis that changes to genes associated with the CO/NCO decision drive divergence in recombination rate between populations. (Cole et al. 2012; Martini et al. 2006).

**Figure 1:**
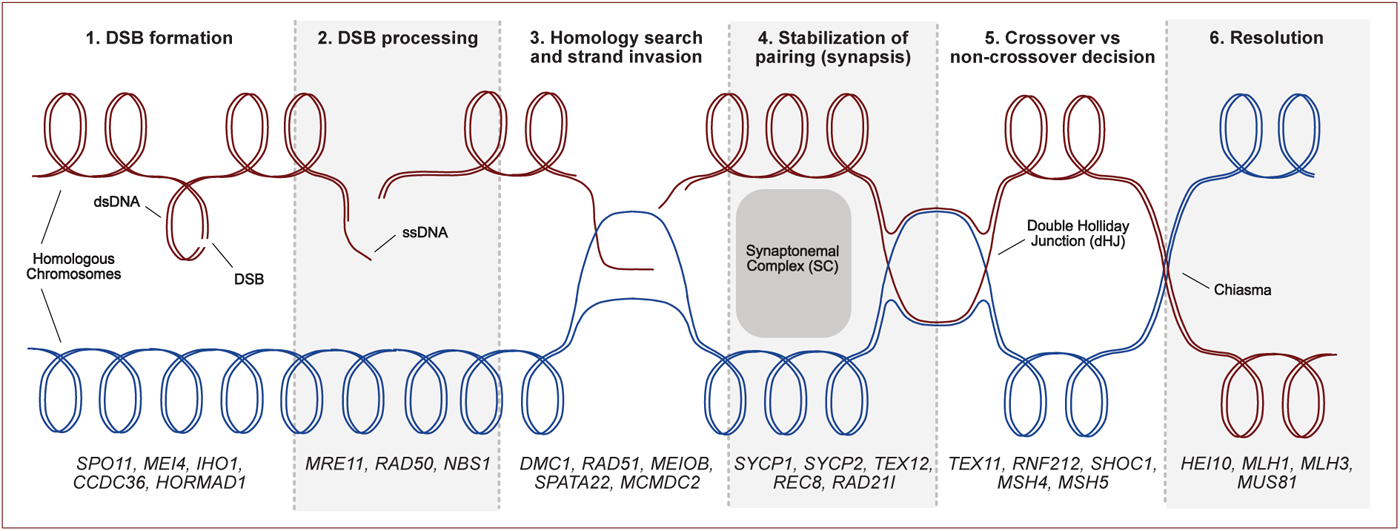
The meiotic recombination pathway can be broken down into six steps. (1) First, double-stranded breaks (DSBs) are non-randomly introduced throughout the genome. (2) Once these breaks are introduced, they are processed via nucleases which leave each DSB with an ssDNA tail. (3) These tails allow DSBs to be lined up appropriately along homologous chromosomes via homology search and strand invasion. (4) The synaptonemal complex (SC) forms a proteinacious scaffold that stabilizes DSBs and homologous chromosomes and creates a structure on which crossover events occur. (5) Though the majority of DSBs are resolved as non-crossovers, a non-randomly distributed subset of the DSBs are designated as crossovers. (6) Finally, DNA repair occurs on both crossovers and non-crossovers, resolving all DSBs. Key genes that regulate each step are listed at the bottom of the figure. Figure modified from Dapper & Payseur 2019.

Birds provide a strong comparative system for investigating patterns of molecular evolution in the meiotic recombination pathway. Both birds and placental mammals have divergence times of *∼* 100 million years (Claramunt and Cracraft 2015; Carlisle et al. 2023; Prum et al. 2015; Kuhl et al. 2021; Liu et al. 2017; Dunwell et al. 2017; Jarvis et al. 2014). Avian diversity is well-represented by an array of high-quality genome assemblies. Interestingly, avian genomes appear to experience recombination rates that are on average twice as high as those observed in mammals (Ellegren 2010; Stapley et al. 2017b) with greater between-species divergence, but higher karyotypic conservation (Zhang et al. 2014; Kapusta et al. 2017; Ellegren 2010). Teleost fish provide an interesting foil to both mammal and avian meiotic recombination pathway evolution. Linkage map length data suggests that fish have both the highest and lowest recombination rates across vertebrates while still maintaining an average recombination rate twice that of mammals (Stapley et al. 2017b). However, it is important to note that teleost fish also have a divergence time over twice that of mammals and birds (Cooney et al. 2021; Takezaki 2018).

Due to their higher recombination rate variability, avian and teleost genomes may afford greater power to identify correlations between molecular evolution and divergence in recombination rates, providing a strong test of the hypothesis that changes to the genes that regulate later stages of the recombination pathway drive divergence in recombination rate between vertebrate genomes. Yet, there are also interesting genomic distinctions between these clades that may result in different selective pressures shaping the evolution of recombination rate and its regulatory pathway. Unlike mammalian and teleost genomes, avian genomes contain numerous microchromosomes, *<*0.5 µm diameter chromosomes that each require a recombination event to segregate properly (Waters et al. 2021; Solovei et al. 1994). Additionally, *PRDM9*, a key, rapidly evolving gene directs recombination hotspots in most mammalian and some teleost genomes (Baker et al. 2017; Paigen and Petkov 2018). But, it is notably absent in avian, yeast, and plant genomes (Paigen and Petkov 2018). These significant differences in recombination mechanism underlie the importance of investigating meiotic recombination rate evolution in non-model, and specifically non-mammalian, organisms.

An important consideration in examining the evolution of the recombination pathway outside of mammals is that most work uncovering the genetics and cellular mechanisms of meiotic recombination has been carried out in model organisms, such as yeast, *Caenorhabditis elegans*, and the house mouse. However, the deep conservation of pathway elements between these distantly related model organisms suggests that we may be able to infer gene function in non-model organisms with some confidence (Bergerat et al. 1997; de Massy 2013; Cole et al. 2010; Zelkowski et al. 2019; Kanaar et al. 1996; Loidl 2016; Kumar et al. 2010; Kan et al. 2011; Hinman et al. 2021).

Here, we use a comparative phylogenetic framework to measure the rates of molecular evolution and identify signatures of positive selection within a panel of key genes in the recombination pathway in avian and teleost clades. Because avian genomes exhibit higher and more variable recombination rates, we predict that avian recombination genes are experiencing elevated rates of molecular evolution and significant signatures of adaptive evolution, as expected of these differences in recombination rate between mammals and birds. Specifically, we expect to see signatures of positive selection within genes driving the CO/NCO decision and synapsis of homologous chromosomes and thus, an association between positively selected genes and genes associated with recombination rate variation. We address the following questions: (1) What patterns of molecular evolution influence recombination rate variation in birds and teleost fish? (2) Do recombination genes evolve more rapidly in non-mammals? (3) Is there more evidence of positive selection in recombination pathway genes in other clades? (4)Are bird and teleost fish recombination genes with signatures of selection also associated with variation in recombination rate? Understanding how patterns of variation in recombination genes differ between clades provides valuable insight into the drivers of recombination rate and thus, the forces governing genetic diversity, selection efficacy, and species divergence.

## Results

### Bird genomes have higher and more variable rates of recombination than mammals

While previous studies have found that avian recombination rates are approximately double those observed in mammals, these comparisons do not control for important karyotypic differences between mammalian and avian genomes (Ellegren 2010; Stapley et al. 2017b). More specifically, avian genomes contain numerous microchromosomes, in addition to macrochromosomes similar to those found in mammalian genomes. Since each microchromosome requires a minimum of one crossover to properly segregate (Ellegren 2010), the extended genetic map length observed in avian genomes could be due to karyotypic differences alone. In other words, the differences in recombination rate could stem from differences in chromosome size and number, rather than differences in the cellular regulation of the number of crossovers. To determine whether recombination rates in avian genomes are higher than than those observed among mammals, when taking karyotype into account, we performed a meta-analysis of published cytological (*MLH1*) studies of recombination rates in avian genomes (Supplementary Data).

We first compared genome-wide measures of recombination rate between avian and mammalian clades, measured in cM/Mb (estimated from *MLH1* data) and in crossovers per haploid chromosomes number (XO/HCN). While birds have significantly higher recombination rates when measured as cM/Mb than mammalian genomes (Figure 2A, *p <* 0.0001, t-test), confirming previous reports, we did not find a significant difference between genome-wide recombination rates between these two clades when measured as XO/HCN (Figure 2B, *p* = 0.1628, t-test). In fact, the trend is towards avian genomes having fewer crossovers per chromosome pair than mammalian genomes. However, both comparisons fail to isolate differences in regulation of recombination rate from karyotypic differences. Just as microchromosomes may increase recombination rate when measured as cM/Mb, they are expected to depress measures of genome-wide recombination when measured as XO/HCN.

To better determine whether avian genomes differ in recombination rate at the chromosomal scale, we compared the average number of *MLH1* foci, a proxy for the number of crossover events, on chromosome 1 - by convention, the largest chromosome in the genome. When measured this way, we found higher and more variable recombination rates among avian genomes (Figure 2C, D, *p* = 0.0108, t-test). Importantly, there is no correlation between the physical size of chromosome 1 and average number of crossovers (Figure 2E, *R*^2^ = 0.0006, *p* = 0.91, linear regression). This is exemplified by comparing largest chromosome in the mouse and chicken genomes. While they are both exceedingly similar in physical size (197 vs. 195 Mbps, respectively), the largest chicken chromosome has on average 9 *MLH1* foci, while the largest mouse chromosome has on average 1.4 *MLH1* foci (Anderson et al. 1999; Nam et al. 2010; Pigozzi 2001). Importantly, heterochiasmy is not implicated in these differences. There was no significant difference in avian sex-specific recombination rate when measured as cM/Mb (*p* = 0.5579), XO/HCN (*p* = 0.1175), or *MLH1* foci on chromosome 1 (*p* = 0.4764)(Supplementary Data). Interestingly, the length of the synaptonemal complex (SC) is considerably longer relative to physical chromosome size in birds than in mammals (*p* = 0.044, paired t-test) and is significantly correlated with recombination rate (Figure 2F, *R^2^* = 0.46, *p* = 0.001, linear regression). This clearly suggests that changes to the proteins in the recombination pathway, specifically those that regulate and/or form the SC, may be responsible for the differences in patterns of recombination between these two clades and motivates a comparative analysis of the molecular evolution of the recombination pathway in these two clades.

**Figure 2:**
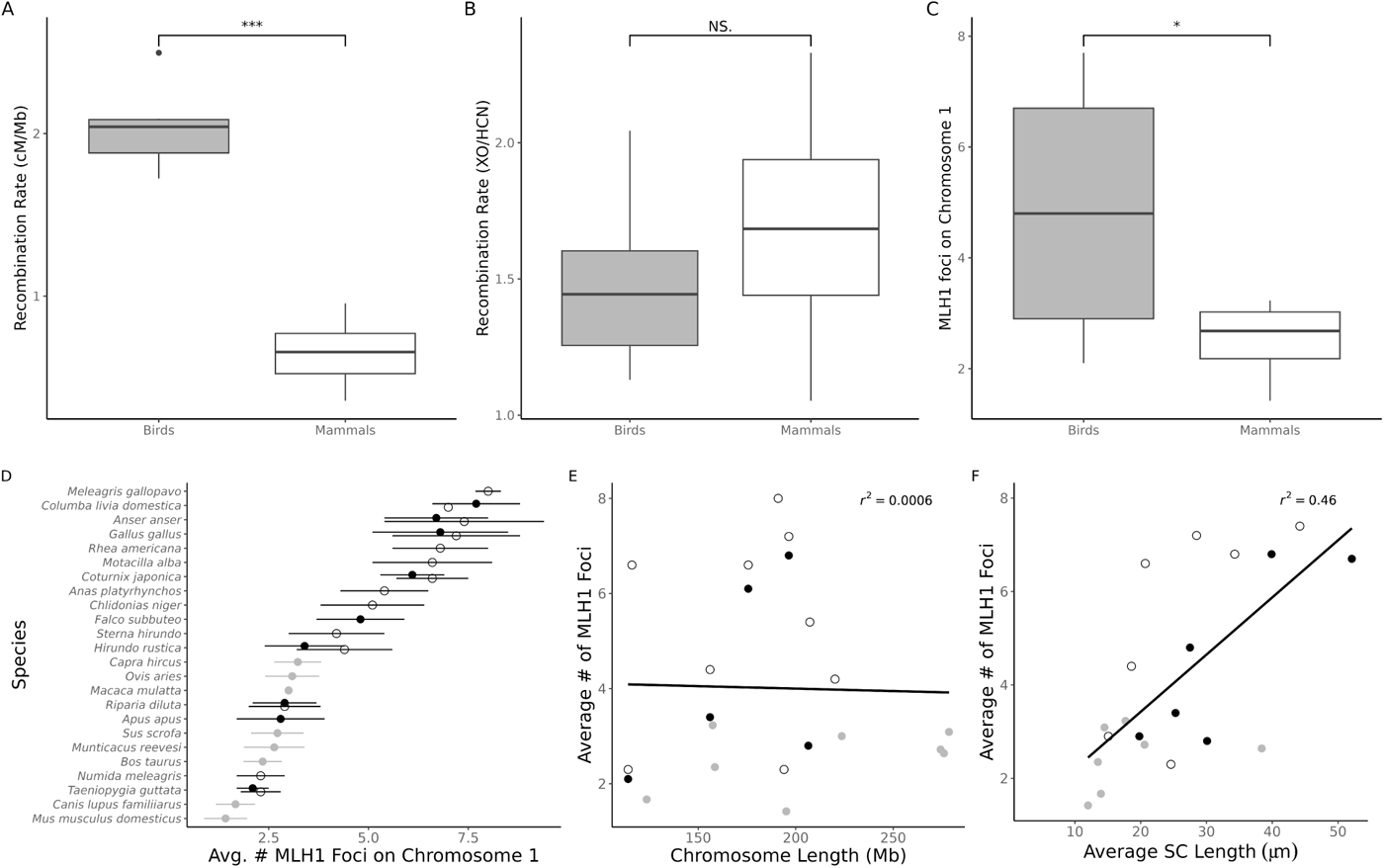
(A) Recombination rate differences between birds and mammals, measured as cM/Mb (*p* < 0.0001). (B) Recombination rate differences between birds and mammals, measured as XO/HCN (crossovers per haploid chromosome number) (*p* = 0.1628). (C) Comparison of number of MLH1 foci on chromosome 1 in birds and mammals (*p* = 0.0108). (D) Species-level comparison of average number of MLH1 foci on chromosome 1 between birds (black) and mammals (grey) by sex, where females are empty circles and males are filled circles. (E) Correlation between length of chromosome 1 and average number of MLH1 foci on chromosome 1 (*R*^2^ = 0.0006, *p* = 0.9115, linear regression). (F) Correlation between average synaptonemal complex length and average number of MLH1 foci on chromosome 1 (*R*^2^ = 0.46, *p* = 0.001, linear regression).

### Phenotypic evolution of meiotic recombination rate differs between clades

In order to contextualize differences in molecular evolution of meiotic recombination genes, it is important to first determine whether phenotypic evolution between clades differs significantly. In order to determine if there is phylogenetic signal within clades, we measured both Blomberg’s K and Pagel’s lambda (Blomberg et al. 2003; Pagel 1999). Notably, there was no significant phylogenetic signal in birds, mammals, or fish for either of these measures (*p >* 0.05, *χ*^2^ test, Data Not Shown). We next compared rates of phenotypic evolution between mammals, birds, and fish with recombination measured as cM/Mb and XO/HCN. We also compared the fit of three models of evolution for each clade: Brownian Motion (BM), Ornstein-Uhlenbeck (OU) and Early Burst (EB). When measured as cM/Mb, there was significantly more support for the multirate model of phenotypic evolution (*σ*^2^_Bird_ = 0.0112, *σ*^2^_Mammal_ = 0.0008, *σ*^2^_Fish_ = 0.0078, *p* = 0.005, *χ*^2^ test). The model of best fit for birds and mammals was BM. There was equal support for both BM and OU models for fish. However, the same trend does not hold when recombination rate is measured as XO/HCN, which controls for large changes in genome size and structure. There was not more support for the multi-rate model over the common rate model (*σ*^2^ _Bird_ = 0.0005, *σ*^2^_Mammal_ = 0.0025, *σ*^2^_Fish_ = 0.0035, *p* = 0.0614, *χ*^2^ test). The model of best fit did not change with the different measure of recombination rate. Thus, we are capturing similar overall levels of phenotypic change in our trait of interest across all three clades.

### Avian and mammalian recombination genes are evolving at similar rates

If the higher and more variable rates of recombination in the avian clade are due to molecular changes in the underlying pathway, we might expect that genes regulating the recombination pathway have evolved more rapidly in the avian clade than the mammalian clade. To test this prediction, we used PAML (Yang 2007) to measure the molecular evolutionary rate, represented by ω, of a panel of 29 key recombination genes across the avian phylogeny (Figure 3A, Supplementary Tables 1, 2) and compared these estimates to those reported for the orthologous genes across the mammalian phylogeny in Dapper and Payseur (2019) (Supplementary Table 3). We found that the evolutionary rates of focal genes across the avian phylogeny ranged from 0.0137 to 0.5778 (mean ω = 0.3123, SD = 0.1606, median = 0.3021 (Figure 4A, Table 1). Within mammals, ω values ranged from 0.0268 to 0.8483 (mean ω = 0.3275, SD = 0.1971, median = 0.3095) (Dapper and Payseur 2019). In comparison with mammals, the average rate of evolution of avian recombination genes is quite similar, and birds show no significant difference in the distribution of ω values (_Bird_ = 0.3123, _Mammal_ = 0.3275, *p* = 0.7718, Wilcoxon signed rank test) (Figure 4A). Comparisons between clades with labeled genes can be found in Supplementary Figures 1-3.

**Figure 3:**
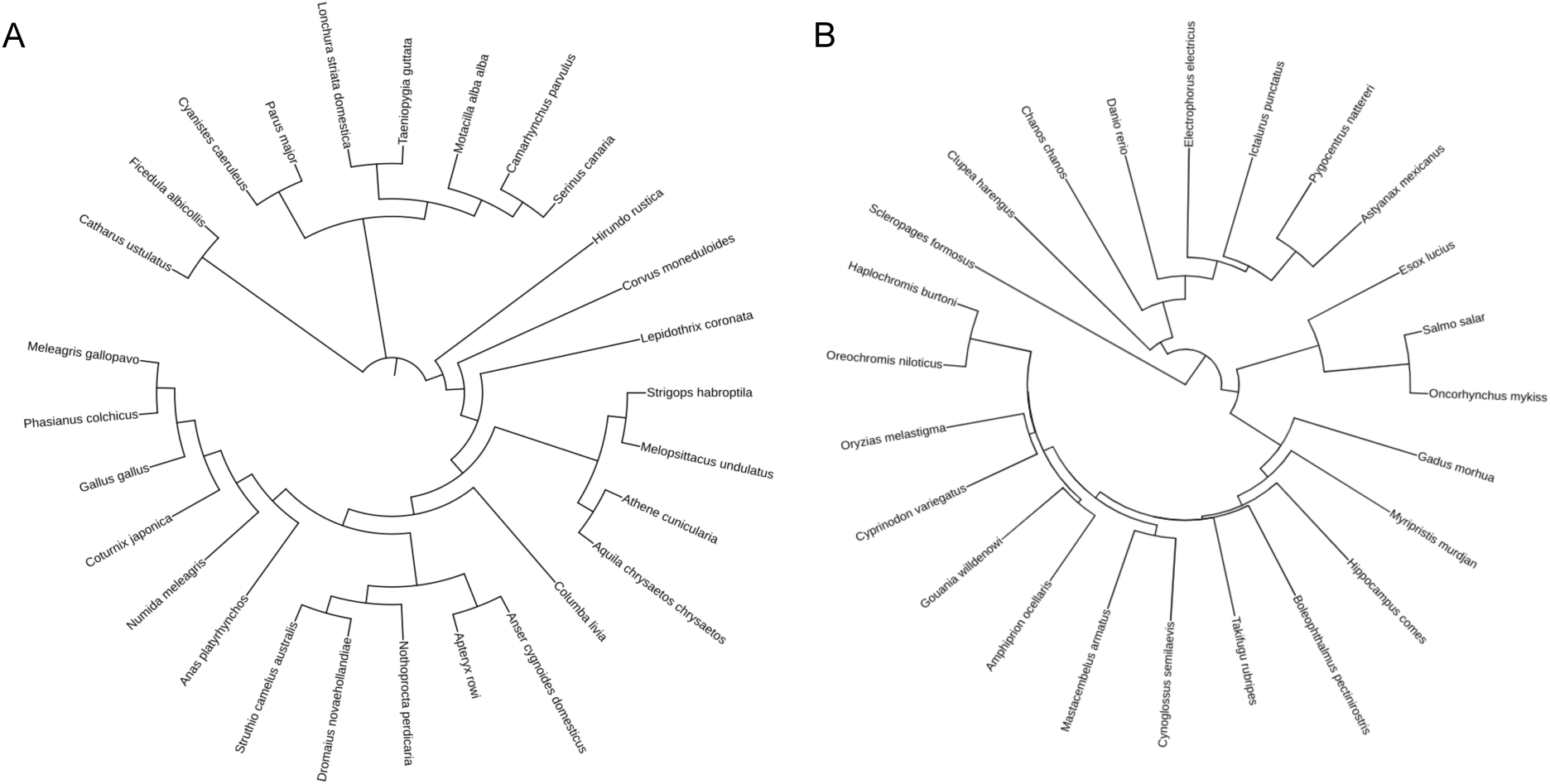
Species tree used for analysis of molecular evolution in bird and teleost fish recombination genes. Teleost fish phylogeny also includes basal teleost *Scleropages formosus*. Figure generated using Letunic and Bork (2021).

**Table 1:**
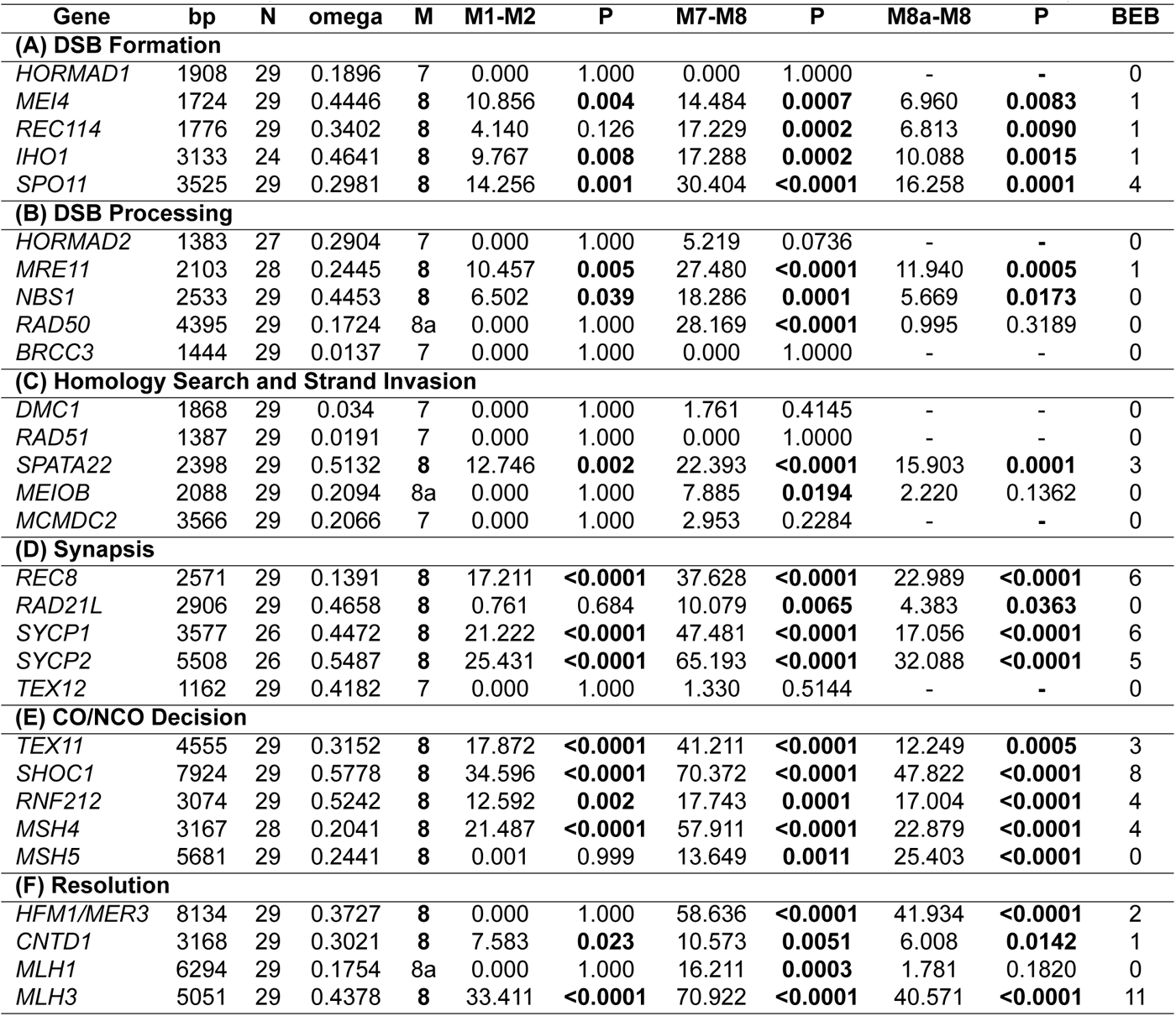
Evolutionary rates and tests for positive selection across birds at 29 recombination genes.

| Gene | bp | N | omega | M | M1-M2 | P | M7-M8 | P | M8a-M8 | P | BEB |
| --- | --- | --- | --- | --- | --- | --- | --- | --- | --- | --- | --- |
| **(A) DSB Formation** |  |  |  |  |  |  |  |  |  |  |  |
| *HORMAD1* | 1908 | 29 | 0.1896 | 7 | 0.000 | 1.000 | 0.000 | 1.0000 | - | - | 0 |
| *MEI4* | 1724 | 29 | 0.4446 | 8 | 10.856 | **0.004** | 14.484 | **0.0007** | 6.960 | **0.0083** | 1 |
| *REC114* | 1776 | 29 | 0.3402 | 8 | 4.140 | 0.126 | 17.229 | **0.0002** | 6.813 | **0.0090** | 1 |
| *IHO1* | 3133 | 24 | 0.4641 | 8 | 9.767 | **0.008** | 17.288 | **0.0002** | 10.088 | **0.0015** | 1 |
| *SPO11* | 3525 | 29 | 0.2981 | 8 | 14.256 | **0.001** | 30.404 | **<0.0001** | 16.258 | **0.0001** | 4 |
| **(B) DSB Processing** |  |  |  |  |  |  |  |  |  |  |  |
| *HORMAD2* | 1383 | 27 | 0.2904 | 7 | 0.000 | 1.000 | 5.219 | 0.0736 | - | - | 0 |
| *MRE11* | 2103 | 28 | 0.2445 | 8 | 10.457 | **0.005** | 27.480 | **<0.0001** | 11.940 | **0.0005** | 1 |
| *NBS1* | 2533 | 29 | 0.4453 | 8 | 6.502 | **0.039** | 18.286 | **0.0001** | 5.669 | **0.0173** | 0 |
| *RAD50* | 4395 | 29 | 0.1724 | 8a | 0.000 | 1.000 | 28.169 | **<0.0001** | 0.995 | 0.3189 | 0 |
| *BRCC3* | 1444 | 29 | 0.0137 | 7 | 0.000 | 1.000 | 0.000 | 1.0000 | - | - | 0 |
| **(C) Homology Search and Strand Invasion** |  |  |  |  |  |  |  |  |  |  |  |
| *DMC1* | 1868 | 29 | 0.034 | 7 | 0.000 | 1.000 | 1.761 | 0.4145 | - | - | 0 |
| *RAD51* | 1387 | 29 | 0.0191 | 7 | 0.000 | 1.000 | 0.000 | 1.0000 | - | - | 0 |
| *SPATA22* | 2398 | 29 | 0.5132 | 8 | 12.746 | **0.002** | 22.393 | **<0.0001** | 15.903 | **0.0001** | 3 |
| *MEIOB* | 2088 | 29 | 0.2094 | 8a | 0.000 | 1.000 | 7.885 | **0.0194** | 2.220 | 0.1362 | 0 |
| *MCMDC2* | 3566 | 29 | 0.2066 | 7 | 0.000 | 1.000 | 2.953 | 0.2284 | - | - | 0 |
| **(D) Synapsis** |  |  |  |  |  |  |  |  |  |  |  |
| *REC8* | 2571 | 29 | 0.1391 | 8 | 17.211 | **<0.0001** | 37.628 | **<0.0001** | 22.989 | **<0.0001** | 6 |
| *RAD21L* | 2906 | 29 | 0.4658 | 8 | 0.761 | 0.684 | 10.079 | **0.0065** | 4.383 | **0.0363** | 0 |
| *SYCP1* | 3577 | 26 | 0.4472 | 8 | 21.222 | **<0.0001** | 47.481 | **<0.0001** | 17.056 | **<0.0001** | 6 |
| *SYCP2* | 5508 | 26 | 0.5487 | 8 | 25.431 | **<0.0001** | 65.193 | **<0.0001** | 32.088 | **<0.0001** | 5 |
| *TEX12* | 1162 | 29 | 0.4182 | 7 | 0.000 | 1.000 | 1.330 | 0.5144 | - | - | 0 |
| **(E) CO/NCO Decision** |  |  |  |  |  |  |  |  |  |  |  |
| *TEX11* | 4555 | 29 | 0.3152 | 8 | 17.872 | **<0.0001** | 41.211 | **<0.0001** | 12.249 | **0.0005** | 3 |
| *SHOC1* | 7924 | 29 | 0.5778 | 8 | 34.596 | **<0.0001** | 70.372 | **<0.0001** | 47.822 | **<0.0001** | 8 |
| *RNF212* | 3074 | 29 | 0.5242 | 8 | 12.592 | **0.002** | 17.743 | **0.0001** | 17.004 | **<0.0001** | 4 |
| *MSH4* | 3167 | 28 | 0.2041 | 8 | 21.487 | **<0.0001** | 57.911 | **<0.0001** | 22.879 | **<0.0001** | 4 |
| *MSH5* | 5681 | 29 | 0.2441 | 8 | 0.001 | 0.999 | 13.649 | **0.0011** | 25.403 | **<0.0001** | 0 |
| **(F) Resolution** |  |  |  |  |  |  |  |  |  |  |  |
| *HFM1/MER3* | 8134 | 29 | 0.3727 | 8 | 0.000 | 1.000 | 58.636 | **<0.0001** | 41.934 | **<0.0001** | 2 |
| *CNTD1* | 3168 | 29 | 0.3021 | 8 | 7.583 | **0.023** | 10.573 | **0.0051** | 6.008 | **0.0142** | 1 |
| *MLH1* | 6294 | 29 | 0.1754 | 8a | 0.000 | 1.000 | 16.211 | **0.0003** | 1.781 | 0.1820 | 0 |
| *MLH3* | 5051 | 29 | 0.4378 | 8 | 33.411 | **<0.0001** | 70.922 | **<0.0001** | 40.571 | **<0.0001** | 11 |

### The teleost fish recombination pathway is more conserved

To determine whether the evolutionary patterns we observe among recombination genes are specific to mammalian and avian clades, we selected teleost fish as outgroup. Teleost, or ray-finned fishes, diverged from the tetrapod lineage between 350-400 million years ago (Near et al. 2012). Notably, teleost fish lack microchromosomes, and thus have a similar chromosomal structure to mammals. Yet, meta-analyses of genetic maps show they have higher and more variable rates of recombination than mammal (when measured as cM/Mb) (Stapley et al. 2017b). Thus, we may predict to also observe similarly elevated rates of evolution of the recombination pathway across the teleost clade. To test this prediction, we again used PAML (Yang 2007) to measure the molecular evolutionary rate, represented by ω, of the panel of 28 key recombination genes across the teleost and basal teleost phylogeny (Figure 3B, Supplementary Table 4). *TEX12* was not consistently identified in teleosts, and was thus left out. In contrast with our prediction, teleost fish recombination genes are evolving significantly more slowly than either bird or mammal recombination genes, with a mean ω value of 0.211 (Figure 4A, Supplementary Tables 5 and 6, *p* = 0.012 and 0.016, respectively, Wilcoxon signed rank test).

### The recombination pathway is experiencing similar selective pressures across vertebrates

**Figure 4:**
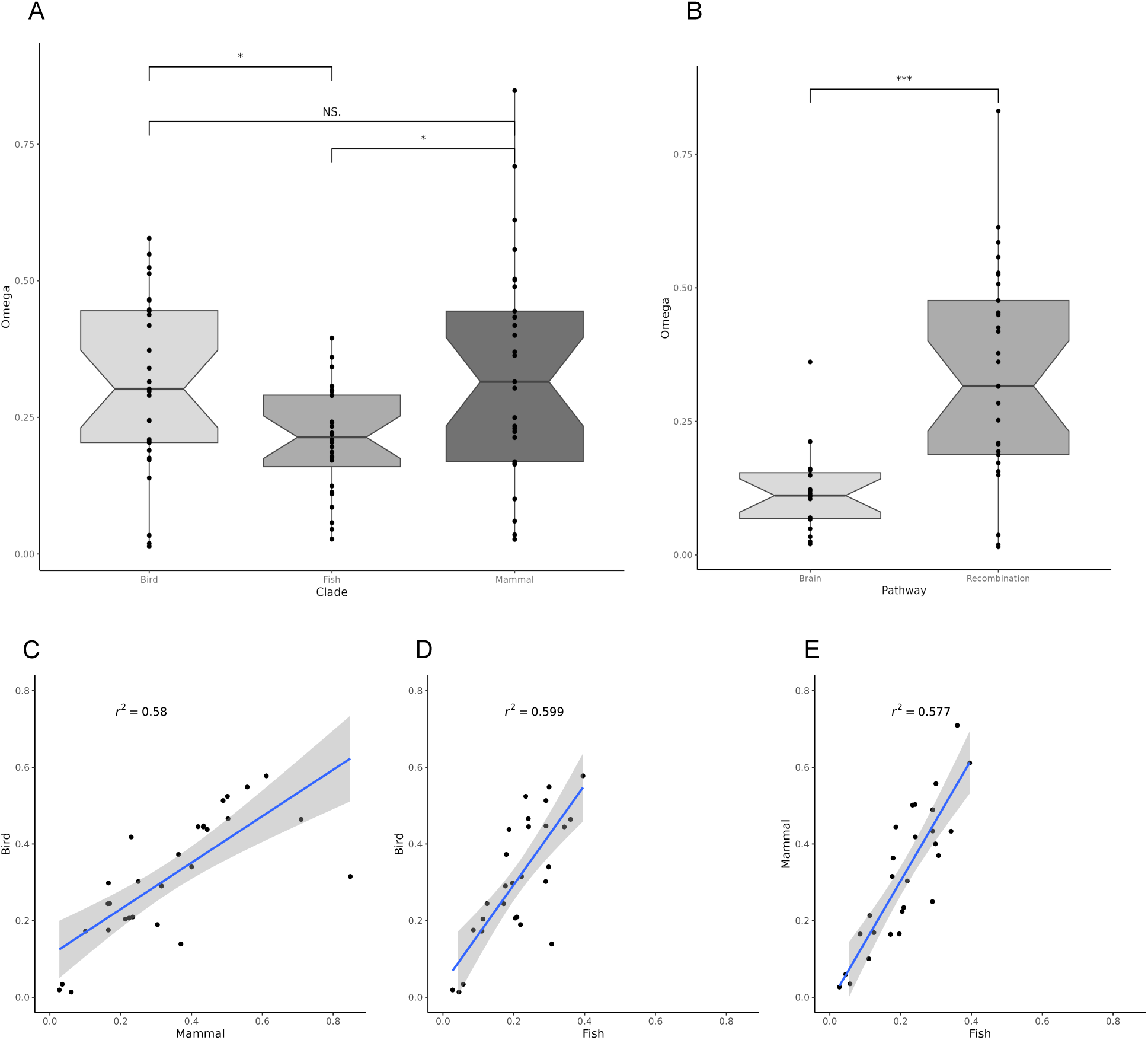
(A) Distribution of evolution rates of recombination genes between birds, teleost fish and mammals. (B) Distribution of evolution rates between genes in the SHH (brain development) and recombination pathways in birds. Correlation of recombination pathway-associated gene omega values between (C) Birds and mammals (R^2^ = 0.58), (D) Birds and teleost fish (R^2^ = 0.599), and (E) Teleost fish and mammals (R^2^ = 0.577). Grey shading represents the 95 % confidence interval for the linear regression.

If the recombination pathway experiences similar selective pressures in vertebrate genomes, we predict that the rate of evolution between orthologous genes will be highly correlated across all three clades. Conversely, striking differences in evolutionary rates between clades may indicate genes, or steps within the pathway, that are experiencing divergent selection pressures. We found that omega (ω) values for each gene are significantly correlated between birds and mammals, supporting the prediction that they have experienced similiar selection pressures (*R*^2^ = 0.58, *p <* 0.0001, linear regression) (Figure 4C). Despite overall lower evolutionary rates of recombination genes in teleosts (Figure 4A), we still observed a significant correlation in ω values between birds and teleosts (*R*^2^ = 0.599, *p* =*<* 0.0001, linear regression) and mammals and teleosts (*R*^2^ = 0.577, *p <* 0.0001, linear regression) (Figure 4D,E), indicating that overall the pathway is experiencing similar selective landscapes. Consistent with this overall trend, there is considerable overlap between the genes evolving the most conserved and most rapidly evolving genes in the pathway between clades (Table 2).

**Table 2:** Genes with elevated (1 SD above the mean) and lower (1 SD below the mean) rates of molecular evolution by clade.

| Gene | Clade |
| --- | --- |
| **Omega 1 SD above mean** |  |
| Mammal | *IHO1, SHOC1, SYCP2, TEX11,* |
| Bird | *RNF212, SPATA22, SYCP2* |
| Fish | *IHO1, MEI4* |
| **Omega 1 SD below mean** |  |
| Mammal | *BRCC3, DMC1, RAD51, RAD50, HEI10* |
| Bird | *DMC1, RAD51, REC8* |
| Fish | *RAD50, RAD51, MLH1* |

**Table 3:** Evolutionary rates and tests for positive selection across birds at 19 brain development genes.

| Gene | bp | N | omega | M | M1-M2 | P | M7-M8 | P | M8a-M8 | P | BEB |
| --- | --- | --- | --- | --- | --- | --- | --- | --- | --- | --- | --- |
| *BMI1* | 3033 | 20 | 0.0205 | 7 | 0 | 1 | 0.006 | 0.9970 | - | - | 0 |
| *CCNA2* | 1917 | 27 | 0.1225 | 7 | 0 | 1 | 3.794 | 0.1501 | - | - | 0 |
| *CCNB1* | 1500 | 13 | 0.1611 | 7 | 0 | 1 | 0 | 1.0000 | - | - | 0 |
| *CCND1* | 4348 | 29 | 0.0341 | 7 | 0 | 1 | 0 | 0.9990 | - | - | 0 |
| *CCND2* | 5663 | 29 | 0.0666 | 7 | 0 | 1 | 0 | 1.0000 | - | - | 0 |
| *DHH* | 3158 | 23 | 0.0693 | 7 | 0 | 1 | 0.944 | 0.6238 | - | - | 0 |
| *EN1* | 2045 | 24 | 0.1204 | 7 | 0 | 1 | 0.413 | 0.8135 | - | - | 0 |
| *EN2* | 2032 | 20 | 0.0490 | 7 | 0 | 1 | 0 | 1.0000 | - | - | 0 |
| *FOXM1* | 3743 | 28 | 0.3612 | **8** | 14.747 | **0.0006** | 37.741 | **<0.0001** | 16.195 | **0.0001** | **1** |
| *GLI1* | 4437 | 28 | 0.2122 | 7 | 0 | 1 | 2.525 | 0.2830 | - | - | 0 |
| *GLI2* | 7564 | 28 | 0.1112 | 8a | 0 | 1 | 7.514 | **0.0234** | 0.828 | 0.3628 | 0 |
| *GLI3* | 9272 | 29 | 0.1586 | **8** | 0 | 1 | 17.750 | **0.0001** | 6.662 | **0.0099** | **6** |
| *IGF2* | 5722 | 29 | 0.0697 | 7 | 0 | 1 | 0 | 1.0000 | - | - | 0 |
| *IHH* | 2679 | 28 | 0.1094 | 7 | 0 | 1 | 3.906 | 0.1418 | - | - | 0 |
| *MYCN* | 1326 | 28 | 0.1595 | 7 | 0.306 | 0.8579 | 2.075 | 0.3543 | - | - | 0 |
| *PTCH1* | 7679 | 27 | 0.1048 | 8a | 0 | 1 | 12.513 | **0.0019** | 1.006 | 0.3159 | 0 |
| *SHH* | 1535 | 28 | 0.1490 | **8** | 13.860 | **0.0010** | 20.561 | **<0.0001** | 12.820 | **0.0003** | **5** |
| *SMO* | 3257 | 27 | 0.1152 | 8a | 5.542 | 0.0626 | 11.972 | **0.0025** | 2.107 | 0.1466 | 0 |
| *SUFU* | 7842 | 29 | 0.0248 | 7 | 0 | 1 | 3.984 | 0.1365 | - | - | 0 |

### Key recombination genes exhibit striking differences in evolutionary rate among clades

Despite the overall concordance between evolutionary rates of recombination genes across the avian and mammalian phylogeny, *TEX11*, a gene whose evolutionary rate is positively correlated with recombination rate in mammals (Dapper and Payseur 2019), shows distinctly different evolutionary patterns in the two clades. While *TEX11* has an ω value of more than two SD above the mean in mammals(ω*_TEX11_*= 0.8483), its evolutionary rate is nearly identical to the mean in birds (ω*_TEX11_* = 0.3152). Notably, *TEX11* is the only recombination gene identified as an influential point through Cook’s distance (Supplemental Figure 4A-4C, D = 2.06, threshold = 0.5) (Cook 1977). As follows, the correlation between avian and mammalian ω values increases considerably with the removal of *TEX11* (*R*^2^ = 0.759, *p <* 0.0001, linear regression). However, *TEX11* is only an influential point in comparisons between birds and mammals.

### The avian recombination pathway exhibits evidence of elevated rates of positive selection

Dapper and Payseur (2019) identified evidence of positive selection in 11 recombination genes across the mammalian phylogeny, significantly concentrated in the steps of the pathway that regulate synapsis and the crossover/non-crossover decision. If the increased variation in recombination rate in the avian clade is adaptive, we predict that we may observe more evidence of positive selection on genes underlying variation in this trait. If the pathway responds to selection in a predictable manner, we also predict that this adaptive evolution will be concentrated in avian recombination genes associated with the same steps of the pathway.

As predicted, we observed significantly more genes exhibiting signatures of positive selection in birds than in mammals (*p* = 0.0214, Fisher’s exact test) (Table 1). More specifically, we identified signatures of positive selection in 19 of 29 (65.5%) recombination genes in birds: *MEI4, REC114, IHO1, SPO11, MRE11, NBS1/NBN, SPATA22, REC8, RAD21L, SYCP1, SYCP2, TEX11, SHOC1, RNF212, MSH4, MSH5, HFM1/MER3, CNTD1,* and *MLH3*. Because multinucleotide mutations (MNMs) can cause false positive signatures of selection (Venkat et al. 2018), we reanalyzed all genes for which Model 8 was the best fit after removing codons that HyPhy’s FitMultiModel reported as potential MNM sites (Lucaci et al. 2021). After this removal, all genes retained their signatures of positive selection except *CNTD1, NBS* and *MSH5* (Supplementary Table 7).

To determine whether the elevated signatures of adaptive evolution we observed was specific to the recombination pathway, or a more general feature of the avian genome, we conducted the same set of analyses on a control panel of genes. We selected a conserved pathway involved with the Sonic Hedgehog (SHH) pathway of brain development (*BMI1, CCNA2, CCNB1, CCND1, CCND2, DHH, EN1,EN2, FOXM1, GLI1, GLI2, GLI3, IGF2, IHH, MYCN, PTCH1, SHH, SMO, SUFU*) (Carballo et al. 2018; Liu et al. 2014; Oliver et al. 2003; Vaillant and Monard 2009). While we did observe evidence of positive selection in genes involved in this pathway across the avian phylogeny, the incidence was not significantly higher than that observed in mammals (Table 3, *p* = 0.6052, Fisher’s exact test). Out of 19 genes, 3 (15.8%) showed evidence of positive selection(Model 8): *FOXM1, GLI3* and *SHH*. We see similar trends in mammals, identifying one gene with signatures of positive selection (*GLI2*, Supplemental Table 8). As expected, there is a significant difference in the ratio of genes with signatures of positive selection between the SHH pathway and the avian meiotic recombination pathway (*p* = 0.001, Fisher’s exact test). Additionally, genes involved in the meiotic recombination pathway evolve at significantly higher rates in birds than those in the SHH pathway (Figure 4B, *p <* 0.0001, Welch’s t-test).

### There is little evidence of positive selection in the teleost recombination pathway

We see a starkly different pattern in teleost fish. Only two recombination pathway genes showed signatures of positive selection within teleost fish: *DMC1* and *RAD50* (Supplementary Table 5). These signatures persist with the removal of possible multi-nucleotide mutation sites. Interestingly, both *DMC1* and *RAD50* are highly conserved despite maintaining signatures of positive selection (ω*_DMC1_* = 0.0572, ω*_RAD50_* = 0.1101), suggesting that these signatures of selection likely come from a small number of sites. Thus, the elevated levels of positive selection in the recombination pathway do not appear to be universal to all vertebrate clades.

### Signatures of positive selection are not concentrated in specific steps of the avian recombination pathway

All genes that had signatures of positive selection in mammals also had signatures of positive selection in birds, resulting in a significant correlation between genes under positive selection between the two clades (*p* = 0.0035, Fisher’s exact test). However, in contrast with the observation in mammals, we did not find that signatures of positive selection are concentrated in genes that regulate the CO/NCO decision in avian genomes (Table 4). This is due to the increased incidence of positive selection in other steps of the avian recombination pathway. We did observe a significantly paucity of genes with signatures of positive selection in the steps of the pathway involved in DSB processing and homology search and strand invasion steps, consistent with strong purifying selection acting on the genes that identify and process DNA damage (*p* = 0.0108, Fisher’s exact test, Table 4).

**Table 4:**
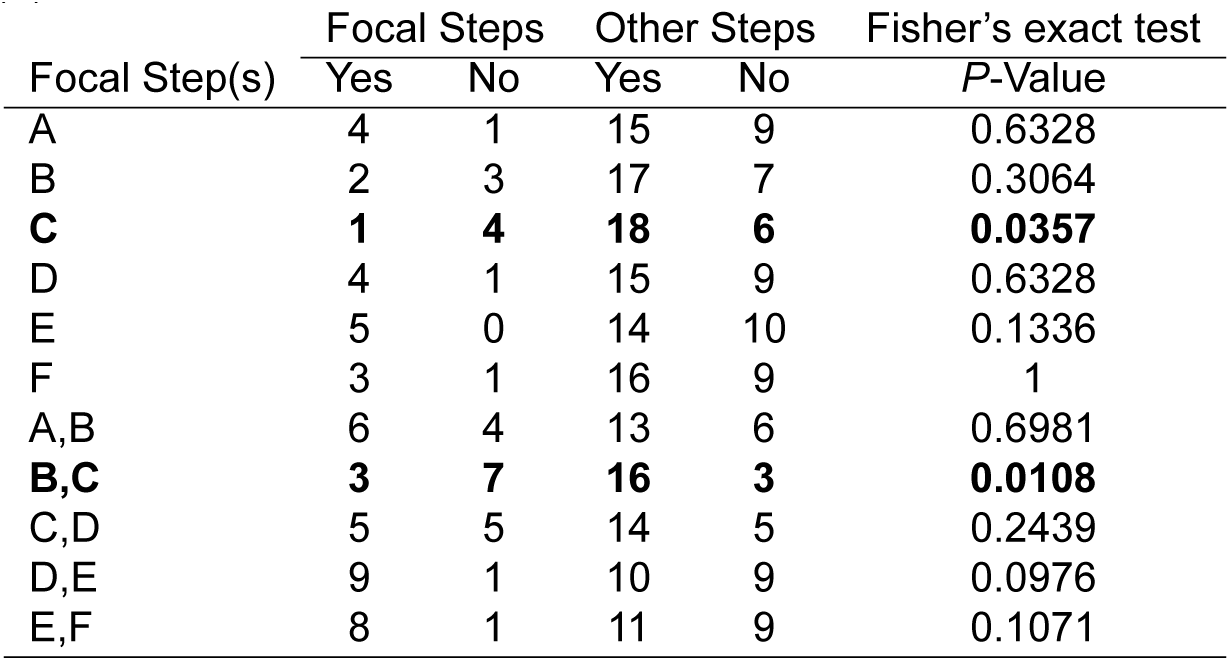
Comparison of proportion of genes with signatures of positive selection by recombination pathway step: (A) DSB formation, (B) DSB processing, (C) Homology search and strand invasion, (D) Synapsis, (E) CO/NCO decision, (F) Resolution.

| Focal Step(s) | Focal Steps |  | Other Steps |  | Fisher's exact test |
| --- | --- | --- | --- | --- | --- |
|  | Yes | No | Yes | No | *P*-Value |
| A | 4 | 1 | 15 | 9 | 0.6328 |
| B | 2 | 3 | 17 | 7 | 0.3064 |
| **C** | **1** | **4** | **18** | **6** | **0.0357** |
| D | 4 | 1 | 15 | 9 | 0.6328 |
| E | 5 | 0 | 14 | 10 | 0.1336 |
| F | 3 | 1 | 16 | 9 | 1 |
| A,B | 6 | 4 | 13 | 6 | 0.6981 |
| **B,C** | **3** | **7** | **16** | **3** | **0.0108** |
| C,D | 5 | 5 | 14 | 5 | 0.2439 |
| D,E | 9 | 1 | 10 | 9 | 0.0976 |
| E,F | 8 | 1 | 11 | 9 | 0.1071 |

### Integrating polymorphism data reveals signatures of purifying selection in the avian recombination pathway

We leveraged polymorphism data within chickens, available for 17 of the recombination genes in our panel, and divergence between chickens and ducks to further explore signatures of selection on the recombination pathway across the avian clade. We identified nine genes with significant McDonald-Kreitman tests, supported by neutrality indices (NI) and direction of selection (DoS) (*p <* 0.05, Fisher’s exact test, Table 5). However, in all cases, the results were consistent with purifying, rather than positive, selection.

**Table 5:** Comparisons of polymorphism within chickens to divergence between chicken and duck at recombination genes.

|  | Pn | Ps | Pn/Ps | Dn | Ds | Dn/Ds | MK Test | alpha | NI | DoS | Direction |
| --- | --- | --- | --- | --- | --- | --- | --- | --- | --- | --- | --- |
| *MEI4* | 26 | 8 | 3.2500 | 81 | 48 | 1.6875 | 0.16 | 1.1368 | 1.9259 | -0.14 | - |
| *REC114* | 6 | 2 | 3.0000 | 53 | 43 | 1.2326 | 0.46 | 1.1979 | 2.4340 | -0.2 | - |
| *IHO1* | 25 | 18 | 1.3889 | 188 | 102 | 1.8431 | 0.4 | 0.9331 | 0.7535 | 0.07 | - |
| *MRE11* | 25 | 6 | 4.1667 | 92 | 117 | 0.7863 | **<0.001** | 1.3663 | 5.2989 | -0.37 | Neg. |
| *NBS1* | 69 | 35 | 1.9714 | 174 | 133 | 1.3083 | 0.09 | 1.0967 | 1.5069 | -0.1 | - |
| *RAD50* | 121 | 51 | 2.3725 | 127 | 284 | 0.4472 | **<0.001** | 1.3945 | 5.3055 | -0.39 | Neg. |
| *BRCC3* | 16 | 6 | 2.6667 | 8 | 72 | 0.1111 | **<0.001** | 1.6273 | 24.0000 | -0.63 | Neg. |
| *RAD51* | 12 | 3 | 4.0000 | 8 | 85 | 0.0941 | **<0.001** | 1.7140 | 42.5000 | -0.71 | Neg. |
| *SPATA22* | 15 | 2 | 7.5000 | 78 | 63 | 1.2381 | **0.01** | 1.3292 | 6.0577 | -0.33 | Neg. |
| *MEIOB* | 85 | 25 | 3.4000 | 44 | 76 | 0.5789 | **<0.001** | 1.4061 | 5.8727 | -0.41 | Neg. |
| *MCMDC2* | 58 | 18 | 3.2222 | 61 | 141 | 0.4326 | **<0.001** | 1.4612 | 7.4481 | -0.46 | Neg. |
| *RAD21L1* | 22 | 8 | 2.7500 | 155 | 122 | 1.2705 | 0.08 | 1.1738 | 2.1645 | -0.17 | - |
| *SYCP1* | 103 | 23 | 4.4783 | 147 | 102 | 1.4412 | **<0.001** | 1.2271 | 3.1074 | -0.23 | Neg. |
| *TEX12* | 1 | 0 | - | 69 | 18 | 3.8333 | - | - | - | - | - |
| *TEX11* | 26 | 9 | 2.8889 | 202 | 51 | 3.9608 | 0.51 | 0.9444 | 0.7294 | 0.06 | - |
| *RNF212* | 11 | 3 | 3.6667 | 15 | 12 | 1.2500 | 0.19 | 1.2302 | 2.9333 | -0.23 | - |
| *HFM1* | 148 | 18 | 8.2222 | 176 | 165 | 1.0667 | **<0.001** | 1.3754 | 7.7083 | -0.38 | Neg. |

### Recombination genes with signatures of positive selection are associated with within-population variation in recombination rate

If the rapid evolution and signatures of positive selection observed among recombination genes are associated with their regulation of recombination rate, we may expect that their rate of molecular evolution to correlated with the rate of evolution of recombination rate. We used Coevol (Lartillot and Poujol 2014) to look for covariation between recombination rate and divergence of recombination genes. Within avians, we used the average number of *MLH1* foci and XO/HCN on chromosome 1 as an estimate of recombination rate. Because of the lack of available *MLH1* foci datasets in teleosts, we estimated XO/HCN from available map lengths. No avian genes showed a correlation between divergence and recombination rate (Table 6, Supplementary Table 6). However, as we were only able to include data from eight species in the avian analysis, our power to detect such correlations is low. Interestingly, we identified significant correlations between divergence in *RAD21L1* and *RNF212* in fish (Table 7). However, this significant correlation does not persist when controlling for variation in the rate of synonymous substitutions (dS).

**Table 6:** Correlations between substitution rate and recombination rate, measured as average MLH1 foci across 8 species of birds for 29 recombination genes. Posterior probabilities are given in parenthesis.

| Gene | Correlation coefficient |  |  | Partial correlation coefficient |  |  |
| --- | --- | --- | --- | --- | --- | --- |
| | dS - $\omega$ | dS - *MLH1* | $\omega$ - *MLH1* | dS - $\omega$ | dS - *MLH1* | $\omega$ - *MLH1* |
| **(A) DSB formation** |  |  |  |  |  |  |
| *HORMAD1* | -0.019 (0.49) | -0.034 (0.47) | -0.507 (0.21) | 0.007 (0.50) | -0.039 (0.47) | -0.369 (0.26) |
| *MEI4* | 0.251 (0.65) | **0.701 (0.96)** | 0.247 (0.65) | 0.170 (0.61) | 0.424 (0.79) | 0.107 (0.57) |
| *REC114* | 0.004 (0.50) | 0.202 (0.61) | 0.259 (0.65) | 0.004 (0.50) | 0.149 (0.60) | 0.189 (0.62) |
| *IHO1* | 0.721 (0.91) | 0.408 (0.85) | 0.368 (0.78) | 0.733 (0.91) | 0.102 (0.57) | 0.090 (0.55) |
| *SPO11* | -0.351 (0.29) | 0.560 (0.91) | -0.308 (0.31) | -0.249 (0.34) | 0.276 (0.70) | -0.176 (0.39) |
| **(B) DSB processing** |  |  |  |  |  |  |
| *HORMAD2* | -0.063 (0.47) | 0.198 (0.61) | 0.052 (0.53) | -0.019 (0.49) | 0.143 (0.59) | 0.047 (0.53) |
| *MRE11* | 0.142 (0.58) | 0.410 (0.73) | 0.364 (0.70) | 0.050 (0.54) | 0.265 (0.68) | 0.267 (0.67) |
| *NBS1* | 0.615 (0.85) | **0.805 (0.96)** | 0.667 (0.88) | 0.286 (0.70) | 0.470 (0.81) | 0.339 (0.72) |
| *RAD50* | -0.468 (0.22) | **0.800 (0.98)** | -0.287 (0.34) | -0.426 (0.22) | 0.677 (0.92) | 0.151 (0.60) |
| *BRCC3* | -0.038 (0.48) | 0.472 (0.76) | -0.022 (0.49) | -0.015 (0.49) | 0.301 (0.70) | -0.005 (0.50) |
| **(C) Homology search and strand invasion** |  |  |  |  |  |  |
| *DMC1* | 0.041 (0.48) | 0.235 (0.63) | -0.058 (0.47) | -0.013 (0.49) | 0.146 (0.60) | -0.041 (0.47) |
| *RAD51* | -0.294 (0.35) | **-0.915 (0.019)** | 0.289 (0.65) | -0.131 (0.41) | -0.605 (0.12) | 0.110 (0.58) |
| *SPATA22* | -0.510 (0.22) | 0.759 (0.94) | -0.499 (0.23) | -0.225 (0.35) | 0.450 (0.80) | -0.231 (0.34) |
| *MEIOB* | -0.170 (0.41) | 0.192 (0.61) | -0.020 (0.49) | -0.068 (0.45) | 0.122 (0.58) | 0.012 (0.50) |
| *MCMD2* | 0.196 (0.62) | 0.533 (0.92) | 0.180 (0.61) | 0.154 (0.59) | 0.272 (0.71) | 0.103 (0.56) |
| **(D) Synapsis** |  |  |  |  |  |  |
| *REC8* | 0.142 (0.59) | -0.021 (0.49) | -0.052 (0.44) | 0.168 (0.60) | -0.015 (0.49) | -0.038 (0.47) |
| *RAD21L1* | 0.342 (0.71) | 0.015 (0.50) | 0.100 (0.57) | 0.413 (0.75) | -0.014 (0.49) | 0.114 (0.57) |
| *SYCP1* | -0.107 (0.44) | -0.122 (0.38) | -0.218 (0.37) | -0.118 (0.43) | -0.092 (0.42) | -0.243 (0.35) |
| *SYCP2* | -0.119 (0.41) | 0.249 (0.69) | -0.309 (0.31) | 0.064 (0.54) | 0.157 (0.61) | -0.298 (0.32) |
| *TEX12* | 0.053 (0.53) | 0.429 (0.81) | 0.138 (0.58) | -0.001 (0.50) | 0.253 (0.69) | 0.115 (0.57) |
| **(E) CO/NCO decision** |  |  |  |  |  |  |
| *TEX11* | 0.092 (0.55) | 0.161 (0.63) | -0.008 (0.50) | 0.132 (0.58) | 0.109 (0.58) | -0.022 (0.49) |
| *SHOC1* | -0.086 (0.44) | 0.082 (0.59) | -0.004 (0.50) | -0.072 (0.46) | 0.063 (0.55) | 0.004 (0.50) |
| *RNF212* | -0.069 (0.46) | 0.078 (0.54) | -0.233 (0.37) | -0.007 (0.49) | 0.050 (0.53) | -0.165 (0.39) |
| *MSH4* | -0.055 (0.46) | 0.318 (0.76) | 0.075 (0.55) | -0.065 (0.45) | 0.206 (0.66) | 0.085 (0.55) |
| *MSH5* | -0.502 (0.20) | 0.226 (0.63) | 0.157 (0.58) | -0.484 (0.22) | 0.153 (0.59) | 0.284 (0.64) |
| **(F) Resolution** |  |  |  |  |  |  |
| *HFM1/MER3* | 0.211 (0.63) | -0.013 (0.48) | -0.029 (0.48) | 0.233 (0.64) | 0.013 (0.51) | -0.044 (0.47) |
| *CNTD1* | -0.082 (0.46) | 0.219 (0.76) | -0.064 (0.47) | -0.021 (0.49) | 0.326 (0.71) | -0.025 (0.48) |
| *MLH1* | 0.129 (0.58) | 0.289 (0.67) | 0.294 (0.69) | 0.094 (0.56) | 0.217 (0.64) | 0.188 (0.63) |
| *MLH3* | 0.306 (0.69) | 0.358 (0.84) | 0.065 (0.56) | 0.349 (0.71) | 0.221 (0.68) | -0.061 (0.46) |

**Table 7:**
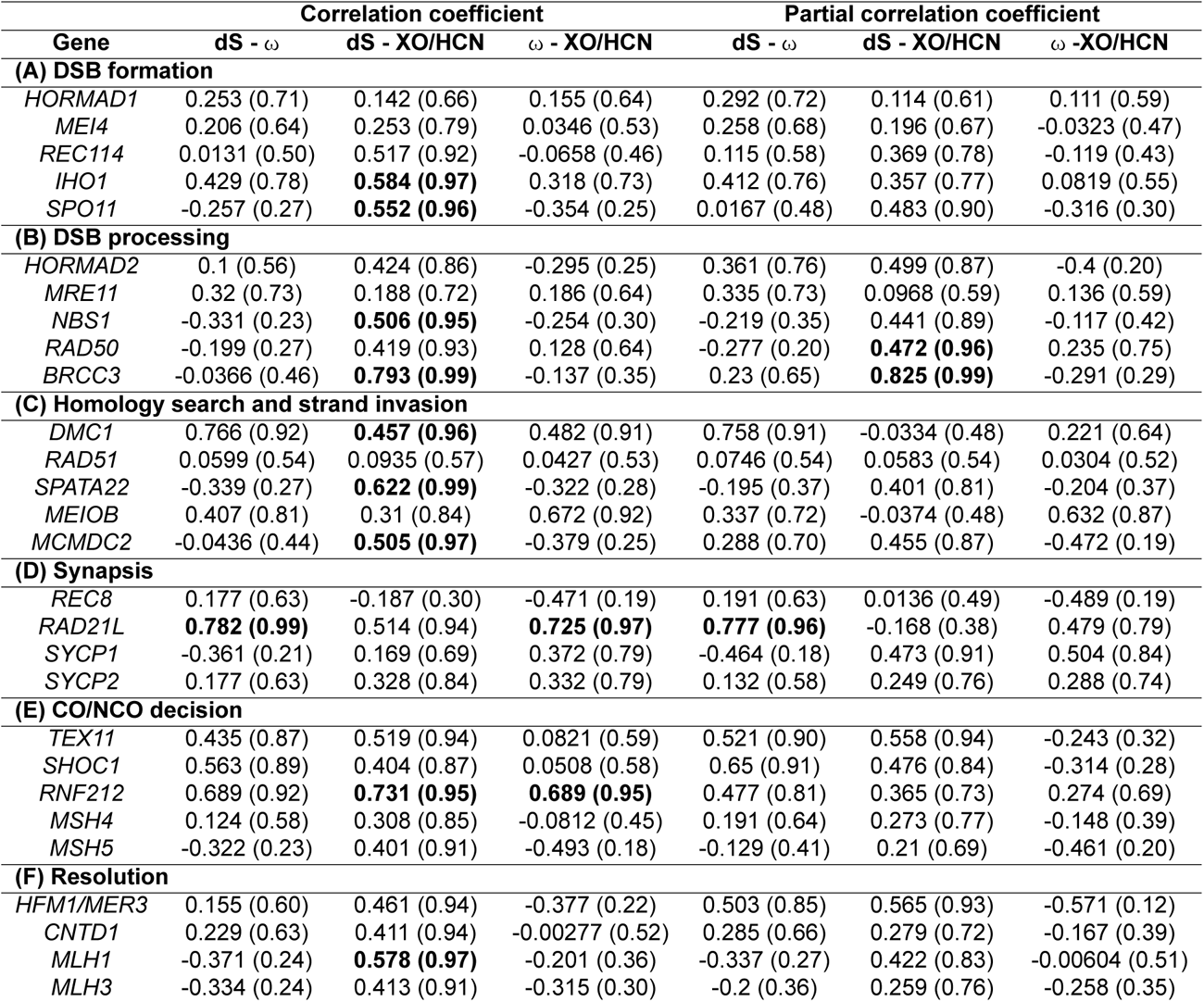
Correlations between substitution rate and recombination rate, measured as XO/HCN across 13 species of fish for 28 recombination genes. Posterior probabilities are given in parenthesis.

| Gene | Correlation coefficient |  |  | Partial correlation coefficient |  |  |
| --- | --- | --- | --- | --- | --- | --- |
| | dS - $\omega$ | dS - XO/HCN | $\omega$ - XO/HCN | dS - $\omega$ | dS - XO/HCN | $\omega$ - XO/HCN |
| **(A) DSB formation** |  |  |  |  |  |  |
| *HORMAD1* | 0.253 (0.71) | 0.142 (0.66) | 0.155 (0.64) | 0.292 (0.72) | 0.114 (0.61) | 0.111 (0.59) |
| *MEI4* | 0.206 (0.64) | 0.253 (0.79) | 0.0346 (0.53) | 0.258 (0.68) | 0.196 (0.67) | -0.0323 (0.47) |
| *REC114* | 0.0131 (0.50) | 0.517 (0.92) | -0.0658 (0.46) | 0.115 (0.58) | 0.369 (0.78) | -0.119 (0.43) |
| *IHO1* | 0.429 (0.78) | **0.584 (0.97)** | 0.318 (0.73) | 0.412 (0.76) | 0.357 (0.77) | 0.0819 (0.55) |
| *SPO11* | -0.257 (0.27) | **0.552 (0.96)** | -0.354 (0.25) | 0.0167 (0.48) | 0.483 (0.90) | -0.316 (0.30) |
| **(B) DSB processing** |  |  |  |  |  |  |
| *HORMAD2* | 0.1 (0.56) | 0.424 (0.86) | -0.295 (0.25) | 0.361 (0.76) | 0.499 (0.87) | -0.4 (0.20) |
| *MRE11* | 0.32 (0.73) | 0.188 (0.72) | 0.186 (0.64) | 0.335 (0.73) | 0.0968 (0.59) | 0.136 (0.59) |
| *NBS1* | -0.331 (0.23) | **0.506 (0.95)** | -0.254 (0.30) | -0.219 (0.35) | 0.441 (0.89) | -0.117 (0.42) |
| *RAD50* | -0.199 (0.27) | 0.419 (0.93) | 0.128 (0.64) | -0.277 (0.20) | **0.472 (0.96)** | 0.235 (0.75) |
| *BRCC3* | -0.0366 (0.46) | **0.793 (0.99)** | -0.137 (0.35) | 0.23 (0.65) | **0.825 (0.99)** | -0.291 (0.29) |
| **(C) Homology search and strand invasion** |  |  |  |  |  |  |
| *DMC1* | 0.766 (0.92) | **0.457 (0.96)** | 0.482 (0.91) | 0.758 (0.91) | -0.0334 (0.48) | 0.221 (0.64) |
| *RAD51* | 0.0599 (0.54) | 0.0935 (0.57) | 0.0427 (0.53) | 0.0746 (0.54) | 0.0583 (0.54) | 0.0304 (0.52) |
| *SPATA22* | -0.339 (0.27) | **0.622 (0.99)** | -0.322 (0.28) | -0.195 (0.37) | 0.401 (0.81) | -0.204 (0.37) |
| *MEIOB* | 0.407 (0.81) | 0.31 (0.84) | 0.672 (0.92) | 0.337 (0.72) | -0.0374 (0.48) | 0.632 (0.87) |
| *MCMD2* | -0.0436 (0.44) | **0.505 (0.97)** | -0.379 (0.25) | 0.288 (0.70) | 0.455 (0.87) | -0.472 (0.19) |
| **(D) Synapsis** |  |  |  |  |  |  |
| *REC8* | 0.177 (0.63) | -0.187 (0.30) | -0.471 (0.19) | 0.191 (0.63) | 0.0136 (0.49) | -0.489 (0.19) |
| *RAD21L* | **0.782 (0.99)** | 0.514 (0.94) | **0.725 (0.97)** | **0.777 (0.96)** | -0.168 (0.38) | 0.479 (0.79) |
| *SYCP1* | -0.361 (0.21) | 0.169 (0.69) | 0.372 (0.79) | -0.464 (0.18) | 0.473 (0.91) | 0.504 (0.84) |
| *SYCP2* | 0.177 (0.63) | 0.328 (0.84) | 0.332 (0.79) | 0.132 (0.58) | 0.249 (0.76) | 0.288 (0.74) |
| **(E) CO/NCO decision** |  |  |  |  |  |  |
| *TEX11* | 0.435 (0.87) | 0.519 (0.94) | 0.0821 (0.59) | 0.521 (0.90) | 0.558 (0.94) | -0.243 (0.32) |
| *SHOC1* | 0.563 (0.89) | 0.404 (0.87) | 0.0508 (0.58) | 0.65 (0.91) | 0.476 (0.84) | -0.314 (0.28) |
| *RNF212* | 0.689 (0.92) | **0.731 (0.95)** | **0.689 (0.95)** | 0.477 (0.81) | 0.365 (0.73) | 0.274 (0.69) |
| *MSH4* | 0.124 (0.58) | 0.308 (0.85) | -0.0812 (0.45) | 0.191 (0.64) | 0.273 (0.77) | -0.148 (0.39) |
| *MSH5* | -0.322 (0.23) | 0.401 (0.91) | -0.493 (0.18) | -0.129 (0.41) | 0.21 (0.69) | -0.461 (0.20) |
| **(F) Resolution** |  |  |  |  |  |  |
| *HFM1/MER3* | 0.155 (0.60) | 0.461 (0.94) | -0.377 (0.22) | 0.503 (0.85) | 0.565 (0.93) | -0.571 (0.12) |
| *CNTD1* | 0.229 (0.63) | 0.411 (0.94) | -0.00277 (0.52) | 0.285 (0.66) | 0.279 (0.72) | -0.167 (0.39) |
| *MLH1* | -0.371 (0.24) | **0.578 (0.97)** | -0.201 (0.36) | -0.337 (0.27) | 0.422 (0.83) | -0.00604 (0.51) |
| *MLH3* | -0.334 (0.24) | 0.413 (0.91) | -0.315 (0.30) | -0.2 (0.36) | 0.259 (0.76) | -0.258 (0.35) |

We also predict that genes associated with within-population variation in recombination rate are more likely to contribute to between species divergence in recombination rate and exhibit signature of positive selection. While there is limited data on genes that are associated with within population variation in avian species, nine recombination genes included in our panel have been associated with variation in recombination rate in mammals: *REC114, REC8, RAD21L. RNF212, TEX11, MSH4, MSH5, HFM1* and *MLH3* (Kong et al. 2008; Sandor et al. 2012; Johnston et al. 2016; Kadri et al. 2016; Petit et al. 2017; Chowdhury et al. 2009; Shen et al. 2018; Ma et al. 2015). Interestingly, all nine recombination genes associated with recombination rate variation in mammals have signatures of positive selection in birds. Compared to other genes in the pathway, recombination genes associated with variation in recombination rate in the mammalian clade are significantly more likely to have signatures of positive selection in the avian clade (*p* = 0.0114, Fisher’s exact test).

## Discussion

Meiotic recombination is a near-ubiquitous biological process, necessary for the production of viable gametes. However, the consequences of meiotic recombination are not relegated only to gamete production; this essential process moderates inter- and intra-genomic genetic variation by generating new combinations of alleles and significantly impacts selection efficacy. Therefore, the rate at which recombination occurs within and between species shapes species divergence (Begun and Aquadro 1992; Brand et al. 2018; Brooks and Marks 1986; Dumont et al. 2009; Kulathinal et al. 2008; Lercher and Hurst 2002; Nachman and Payseur 2012; Ptak et al. 2005; Tigano et al. 2022). Importantly, recombination rate itself evolves and exhibits striking patterns of divergence between populations, even over relatively short time scales (Dumont et al. 2009; Johnston et al. 2016; Kadri et al. 2016; Petit et al. 2017; Johnston et al. 2018; Sandor et al. 2012; Kong et al. 2008). Recombination events arise through a highly regulated cellular pathway (Youds and Boulton 2011; Cannavo et al. 2020; Baudat et al. 2013; Keeney 2001), indicating that molecular changes in the underlying genes shape the phenotypic evolution of this fundamental genomic parameter. This observation raises the prospect that a comparative study of the molecular evolution of the genes that regulate this pathway can reveal the genetic architecture of divergence in recombination rate.

We observe highly correlated rates of molecular evolution within the meiotic recombination pathway across three vertebrate clades. This result suggests that the evolutionary pressures (i.e. selection, drift, and mutation) that shape recombination rate are largely consistent across vertebrates. Given the overall consistency of the rate of evolution of recombination genes, significant outliers may identify clade-specific drivers of the evolution of recombination rate. Within our panel of key recombination genes, we identified one notable outlier, *TEX11*, in the comparison of evolutionary rates between the avian and mammalian clades. While *TEX11* evolves very rapidly in mammals (Dapper and Payseur 2019), its evolutionary rate across birds and teleost fish is considerably slower and on par with the average rate of evolution of the recombination pathway in these clades. *TEX11* is particularly interesting because it plays a crucial role in crossover formation and synapsis maintenance (Yang et al. 2008) and its evolutionary rate is correlated with genome-wide recombination rate across mammals (Dapper and Payseur 2019). Notably, all studies of *TEX11* phenotype are from mammalian systems (Ji et al. 2021; Kitayama et al. 2022; Song et al. 2023; Tang et al. 2011; Yang et al. 2008, 2015; Yatsenko Alexander N. et al. 2015; Yu et al. 2021, 2012). Together these observations suggest that *TEX11* may be a mammal-specific driver of recombination rate variation. Thus, despite the overall concordance between evolutionary patterns, clade-specific drivers of the evolution of recombination rate may play important roles in driving divergence.

If recurrent adaptive evolution contributes to divergence in recombination rate, we may expect to observe more signatures of positive selection in clades with higher rates of phenotypic evolution. While differences in genome architecture makes it difficult to disentangle the contribution of karyotypic evolution and molecular evolution to divergence in recombination rate, we leveraged existing *MLH1* data to compare rates of recombination between avian and mammalian genomes. We found that the number of crossovers (*MLH1* foci) on chromosome 1, by convention the largest chromosome in the genome, is higher and more variable in avian genomes. Importantly, the divergence in foci number is not explained by differences in chromosome size, but is positively correlated with the length of the synaptonemal complexes (SCs), the proteinaceous structure that stabilizes the pairing of homologous chromosomes and provides a substrate for the resolution of crossovers, which is longer relative to chromosome size in avian genomes than in mammalian genomes.Thus, these differences are likely driven by molecular changes in the recombination pathway, rather than as a direct result of changes to chromosome size and structure in avian genomes. However, the evolution of microchromosomes in avian genomes may have indirectly generated selection for higher recombination rates in avian genomes to ensure the minimum of one crossover per chromosome necessary for proper chromosomal segregation (Ellegren 2010; Hassold and Hunt 2001).

Consistent with this prediction, we observed a striking elevation in the signatures of positive selection among genes in the recombination pathway across the avian phylogeny, with 19 out of 29 genes surveyed exhibiting significant signatures of selection. This is significantly higher than the 11 out of 32 genes identified with signatures of positive selection in the mammalian recombination pathway (Dapper and Payseur 2019). To account for the possibility of poorer genome quality, we excluded available avian genomes with significant issues in gene alignment prior to data collection and analysis. As maximum likelihood models are prone to false positives, we took a number of steps to verify these results, including hand curation to minimize misalignment, utilization of models that account for multi-nucleotide mutations (Venkat et al. 2018; Lucaci et al. 2021), and reanalysis of the mammalian data to ensure consistency in data preparation and analysis. While in a few cases these measures removed signals of positive selection, there was no meaningful change to the overall result. We also selected a control pathway (SSH pathway) not involved in meiotic recombination, or reproductive processes, which we analyzed using the same pipeline. Our control pathway did not exhibit patterns of rapid evolution, such as that observed in some recombination genes, nor did we observe a significant elevation in incidence of positive selection. Thus, our result is unlikely to be the result of lower overall genome quality among the included avian species.

Importantly, while patterns of molecular evolution can be indicative of recurrent positive selection acting on protein-coding genes, they do not reveal the source of the selective pressure. Thus, while this observation is consistent with the hypothesis that selection favored elevated rates of recombination across the avian phylogeny, it is also possible that these signatures arise as result of pleiotropic selection. If the molecular evolution of these genes is driven by their affect on recombination rate, we may expect to observe: (1) correlations between the rates of phenotypic and molecular evolution, such as the positive correlation between the rate of molecular evolution in *TEX11* and recombination rate across the mammalian phylogeny Dapper and Payseur (2019) and (2) genes in the recombination pathway that influence recombination rate are more likely to exhibit signatures of positive selection than those that do not.

In contrast to this prediction, we did not observe a significant correlation between evolutionary rate and recombination rate across the avian phylogeny. However, it is important to note because we were only able to include eight species in this analysis, we have limited power to detect such a correlation. This lack of correlation may also be consistent with a realistic scenario in which the interplay between genetic drift and purifying selection shape the overall rate of evolution of the gene, and positive selection acted on small subset of impactful sites. Interestingly, we did observe a significant positive correlation between genes associated with within population variation in recombination rate in mammalian populations, and thus likely to impact recombination rate in avian genomes, and those that exhibited a significant signature of positive selection across the avian phylogeny. Given the overall conservation of function of genes in the recombination pathway, this correlation suggests that molecular evolution of these genes across the avian phylogeny may contribute to variation in recombination rate. A more direct comparison did not find a significant correlation with genes that exhibit signatures of positive selection in mammals. However, the overall lower number of genes experiencing positive selection in mammals may reduce our power to identify such a correlation, as there is also a significant correlation between the genes experiencing positive selection in avian and mammalian genomes. All genes that had signatures of positive selection in mammals also had signatures of positive selection in birds, providing additional support for the hypothesis that positive selection may act on the meiotic recombination pathway in predictable ways.

The structure of the recombination pathway suggests that divergence in rate may arise from upstream changes that alter the number of potential crossover sites (DSB formation) and/or downstream changes that regulate the number of potential sites that are resolved as mature crossovers (CO/NCO decision) (Cole et al. 2012; Martini et al. 2006; Johnston et al. 2018; Sandor et al. 2012; Kadri et al. 2016; Petit et al. 2017; Ortiz-Barrientos et al. 2016). Molecular genetic and evolutionary analyses in mammals suggest that protein-coding and/or regulatory changes to genes that regulate the crossover (CO) vs. noncrossover (NCO) decision are most likely to affect the evolution of recombination rate (Dapper and Payseur 2019; Segura et al. 2013; Dumont et al. 2009; Romanienko and Camerini-Otero 2000; Kumar et al. 2010; Yang et al. 2008; Guiraldelli et al. 2018; Reynolds et al. 2013; Kadri et al. 2016; de Vries et al. 1999). However, as a result of the high incidence of signatures of positive selection across the avian meiotic recombination pathway, genes experiencing signatures of positive selection are not concentrated in the CO/NCO decision pathway steps, as seen in mammals. Instead, there are significant signatures of positive selection within the steps of the pathway responsible for the formation of double-stranded breaks, synapsis, and the CO/NCO decision. This supports the hypothesis that changes in proteins that govern SC development and maintenance within the avian genome may contribute to differences in recombination rate between mammals and birds, but also raises the prospect that variation may also be driven by differences in the number of double-strand breaks generated early in the pathway. Conversely, there is evidence of significant purifying selection acting on avian recombination genes that process and identify DNA damage, a trend also seen within mammals. These repeated patterns of positive and purifying selection within recombination pathway genes between clades suggest that while there are significant phenotypic differences between clades, there are predictable effects of selection acting on the meiotic recombination pathway.

Although the rates of evolution of the recombination genes in teleost fish are correlated with their orthologs in mammalian and avian genomes, the pathway is significantly more conserved without widespread evidence of positive selection (only 2 of 28 recombination genes exhibit signatures of positive selection in the teleost clade). One possibility is that due to the longer divergence times, the overall evolutionary rate of recombination rate is lower in this clade. However, the longer divergence time for teleost fish does not seem to explain differences in recombination rate: In the cM/Mb multi-rate models, their *σ*^2^ value was similar to that of birds, and there was not support for the multi-rate model when measuring recombination as XO/HCN. Thus, it appears that recombination rate evolution in teleost fish is not associated with rapid or adaptive evolution of underlying pathway in the same manner as birds and fish.

Understanding the impact of changes to proteins involved in meiotic recombination is crucial to elucidating mechanisms responsible for between-species divergence. This study substantiates the existence of similar selective landscapes acting upon the meiotic recombination pathway between vertebrate clades. Specifically, we identify strikingly similar rates of molecular evolution within key avian and mammalian recombination genes. *TEX11*, an important element of the CO/NCO decision, is a significant outlier to this trend, suggesting the possibility of clade-specific drivers of meiotic recombination rate variation. Additionally, we identify strong between-clade patterns of selection acting on recombination pathway genes: Patterns of positive selection acting on genes mediating recombination rate variation and those responsible for the formation and resolution of double-stranded breaks, and patterns of purifying selection acting upon genes responsible for DNA damage identification and processing, suggesting that changes in recombination pathway proteins drive recombination rate variation.

## Methods

### Meta-Analysis of Cytological Data

We collected data from published cytological studies of avian (N = 9, male; N = 14, female) and mammalian (N = 11, male) genomes (Supplemental Table). All studies inclued in the meta-analysis used immunofluorescence to label MLH1 foci, with two exceptions which used phosphotungstic acid. Due to difference in the feasibility of gamete collection, the mammalian studies almost exclusively report MLH1 data from male sperm cells, while the majority of avian studies report MLH1 data from female egg cells. As we did not observe significant heterochiasmy in avian genomes (*p* = 0.4764, t.test, Supplementary Data), we report data that maximized sample size by comparing female avian MLH1 estimates and male mammalian MLH1 estimates. Analyses that included only male estimates supported the same conclusions and can be found in the supplementary material. In cases where multiple MLH1 foci counts were reported for a single species, we chose to include data from the study with the largest sample size (determined by the number of individuals and total cell count). All included studies reported the average genome-wide MLH1 foci count (a proxy for the total number of crossovers in the genome). We compared two measures of genome-wide recombination rate: (1) total number of *MLH1* foci per haploid chromosome number (XO/HCN) and (2) estimated centiMorgans per Megabase (cM/Mb). To estimate genetic map length from MLH1 foci data, we multiplied the number of *MLH1* foci by 50 cM/foci, giving us an estimate of genetic map length (cM). We then divided total map length by genome size measured in Mb. To estimate genome size using two approaches: (1) by using the total size of the genome assembly (Mb), which may underestimate length because they may not include regions of the genome that are difficult to assemble, and (2) by converting genome weight (C-value, pg) to an estimated genome size (Mb). Genome weights were found using the Animal Genome Size Database (Gregory 2024). For the conversions, we assumed that 1 pg is roughly 978 Mb (Gregory 2024). Both methods of estimating total genome size produced qualitatively similar results (Supplementary Data).

All avian studies and a subset of mammalian studies (N = 8, mammals) reported the average number of *MLH1* foci counts for the largest chromosome in the genome (by convention chromosome 1). To test whether variation in number of *MLH1* foci on Chromosome 1 could be explained by variation in chromosome size, we used the size of Chromosome 1 (Mb) as reported in the lastest genome assembly of each species. Many of the cytological studies we surveyed also reported estimated of the length of the synaptonemal complex on Chromosome 1 (N = 9, male, mammal; N = 6, male, avian; N = 7, female, avian). All analyses were performed using R Statistical Software (v4.3.0; R Core Team 2021R Core Team (2023)).

### Data Acquisition and Processing

We utilized the same panel of 32 genes from Dapper and Payseur (2019) to compare meiotic recombination rate pathway evolution between mammals, birds, and teleosts (Supplementary Table 1). Reference sequences were downloaded from NCBI from 29 bird species and 24 teleost fish species (Figure 3). These species were chosen due to their divergence times and quality of reference sequences. However, we were unable to identify orthologs of all 32 recombination genes in avian and teleost genomes. Thus, 29 candidate genes were investigated in birds (Supplementary Table 1). *RNF212B, HEI10* and *MUS81* were excluded entirely from both birds and teleosts due to a lack of useful data. *TEX12* was not available for teleost fish, so we excluded this gene during analysis, resulting in 28 focal genes for the teleost recombination pathway. Phylogenetic trees were inferred using TimeTree, which allows the generation of trees from species lists (Kumar et al. 2022). We then generated visual trees with iTOL v.5 (Letunic and Bork 2021). For 26 genes, sequences were available from all species of birds. Sequences were not available for one or more individuals for the following genes: *MRE11, SYCP1*, and *MSH4*. Sequence availability proved more difficult in teleost fish. Data was unavailable for one or more teleost fish for 11 genes; however, all genes surveyed had at least 18 teleost species represented.

To control for differences in genome quality between clades in our subsequent analyses, we also selected a focal panel of control genes involved in a brain development pathway, (*BMI1, CCNA2, CCNB1, CCND1, CCND2, DHH, EN1, FOXM1, GLI1, GLI2, GLI3, IGF2, IHH, MYCN, PTCH1, SHH, SMO, SUFU*) (Carballo et al. 2018; Liu et al. 2014; Oliver et al. 2003; Vaillant and Monard 2009). We selected these genes for their inclusion in a well-described pathway and because we expect them to be conserved.

### Measuring Rates of Phenotypic Evolution

In order to compare rates of evolution of recombination rate between our three clades, we needed to identify consistent units of measurement of recombination rate. This was complicated because recombination rate in fish is generally estimated via genetic map length, whereas we have measurements of the number of crossovers per genome in mammals in birds, via MLH1 foci. We compared recombination rate as cM/Mb, by estimating this measure from MLH1 data in birds and mammals, and as XO/HCN, by estimating this measure from genetic map length in fish. We first collected available map lengths for teleost species and estimated recombination rate as cM/Mb by dividing map length by assembly size (Supplementary Data). To estimate genetic map length from MLH1 foci data, we multiplied the number of *MLH1* foci by 50 cM/foci, giving us an estimate of genetic map length (cM) and divided total map length by genome size measured in Mb. In order to estimate recombination rate as XO/HCN for teleost fish, we divided map length by 50 to get estimated crossover number, then divided crossovers by either haploid chromosome number or number of linkage groups. For birds and mammals, we simply divided the number of MLH1 foci by HCN. For both measures of recombination rate, we used the geiger package in R to fit models against the bird, fish and mammal phylogenies (Pennell et al. 2014). For each clade, we fit models of Brownian Motion (BM), Ornstein-Uhlenbeck (OU) and Early Burst (EB) using the ‘fitContinuous()’ function. We then calculated AIC scores for each model and ranked the models using Akaike weights. To determine whether there is more evidence for a multi-rate or common-rate model between phylogenies, we used the ratebytree function from the phytools R package (Revell 2024). We also used the phylosig function from phytools to identify phylo-genetic signal using both Blomberg’s K and Pagel’s lambda (Pagel 1999; Blomberg et al. 2003).

### Phylogenetic Comparative Approach

We used maximum likelihood approaches to assay evidence of selection (using the same approach as (Dapper and Payseur 2019)). We used CODEML site models from PAML to measure the rate of synonymous to non-synonymous substitutions (Yang 2007). CODEML requires a phylogenetic tree, for which we used Newick format, and multiple sequence alignments, for which we used PHYLIP format (Felsenstein 2005). To ensure high quality files for analysis, we aligned reference sequences acquired from NCBI using MUS-CLE and GBlocks via TranslatorX (Edgar 2004; Abascal et al. 2010; Castresana 2000), hand-cleaning when necessary. We aligned bird sequences with their mammalian counterparts to ensure that equivalent regions of the genes were compared. Specific assemblies used may be found in Supplementary Tables 2 and 4.

We used the JC69 model of codon substitution and set the following initial parameters: kappa = 2, omega = 0.4, alpha = 0. Rates were allowed to vary between branches (clock = 0). We calculated these rates for six site models(M0, M1a, M2a, M7, M8 M8a) and determined the model of best fit via likelihood ratio test.

### Identifying Signatures of Selection

A common method used to identify signatures of selection is comparing the ratio of synonymous (dS) to non-synonymous substitutions (dN) (Anisimova et al. 2001). This is represented by = *dN* /*dS*. If there are a larger number of non-synonymous amino acid changes than synonymous changes (higher dN), ω increases. If ω is significantly greater than 1, there is evidence of adaptive evolution. An ω value significantly less than 1 is indicative of purifying selection. Neutrally evolving sites will have an ω value close to 1. PAML’s CODEML utilizes several evolutionary models to calculate the best fit for the data.

To determine model of best fit for bird recombination genes, we first compared Model 1 versus Model 2. M1 assumes that there are two classes of sites: one where ω < 1, and one where ω = 1, indicating neutral evolution. M2 introduces a third class, wherein ω > 1, which indicates positive selection. Models 7 and 8 control for variation in ω across a beta distribution of 0-1. M7 contains 10 of these site classes where ω < 1, whereas M8 includes a class of ω > 1 to allow for positive selection. We compared M7, M8, and an additional model designated M8a. M8a allows ω = 1. This model is frequently a better fit in genes where there are many sites that are evolving neutrally. We also report the number of codons undergoing positive selection (Bayes empirical Bayes, BEB; P > 0.95).

### Multi-Nucleotide Mutations

Multinucleotide mutations (MNMs) occur when two or more mutations in proximity on one haplotype occur during the same mutational event (Wang et al. 2020). These mutations violate the assumptions of many maximum liklihood models of molecular evolution and can potentially contribute to false inferences of positive selection (Venkat et al. 2018; Yang 2007). In order to identify potential MNMs in our dataset, we used the FitMultiModelHyPhy analysis file to identify sites where a simultaneous nucleotide substitution model may fit better (Lucaci et al. 2021). We removed these sites and re-ran CODEML on the edited sequences. This process was done only for genes that showed evidence of positive selection to determine whether this signal remained after the removal of potential MNMs.

### Polymorphism and Divergence

Polymorphism data were acquired from whole genome sequences of Nandao chickens (Yang et al. 2021). The data are available through the European Variant Archive, under the study identifier PRJEB46210. We used the McDonald-Kreitman test with Jukes-Cantor correction to compare synonymous and nonsynonymous substitutions and polymorphisms (Jukes and Cantor 1969; Mcdonald and Kreitman 1991; Haller and Messer 2017). We also measured the neutrality index (NI), which measures divergence from the neutral expectation (pN/pS = dN/dS), and the direction of selection for each gene (Stoletzki and Eyre-Walker 2011).

### Evolutionary Rate Covariance

We used Coevol to identify any correlations between recombination rate and divergence of recombination genes between species. Briefly, Coevol uses Bayesian Markov Chain Monte Carlo (MCMC) methods to estimate correlations between traits and substitution rates in sequences given a phylogenetic tree (Lartillot and Poujol 2014). Instead of linkage map lengths, we used the number of *MLH1* foci on chromosome 1 for eight species from our focal avian species list (*C. japonica, T. guttata, H. rustica, G. gallus, A. platyrhynchos, N. meleagris, M. alba* and *M. gallopavo*) as a stand-in for recombination rate. We used the same parameters as Dapper and Payseur (2019) in Coevol (burn-in = 1,000; MCMC chain = 25,000; relative difference *ω* < 0.01) which returned pairwise correlation coefficients between recombination rate, ω, and dS, along with partial correlation coefficients for each pairwise correlation (Dapper and Payseur 2019). We also investigated correlations between divergence and recombination rate measured as XO/HCN within the same focal bird panel, excluding *M. alba*, and in 13 species from our teleost fish phylogeny (*E. lucius, O. mykiss, S. salar, T. rubripes, C. semilaevis, H. burtoni, O. melastigma, G. morhua, I. punctatus, A. mexicanus, D. rerio, C. harengus* and *S. formosus*).

## Acknowledgements

We thank Austin Drury, Jean-Francois Gout, Federico Hoffmann, Mark Welch, and members of the Dapper and Gout labs for constructive feedback on study design and interpretation of results. This work was supported by NSF CAREER 2143063 to ALD.

## Author Contributions

TSG lead data collection, data analysis and writing. TSG and ALD collaborated on study design, data analysis, and writing. KS, VV, and LZ contributed to data collection and analysis. AKL assisted with identification of the control pathway.

## Data Availability Statement

Raw data may be found in the following repository: https://github.com/tszaszgreen/Meiotic_Recombination

