## Supplementary Tables and Figures for "Molecular evolution of the meiotic recombination pathway in vertebrates"

### Supplementary Material

Szasz-Green et al. 2024

June 27, 2024

**Supplementary Tables**

Supplementary Table 1: List of 29 genes surveyed, organized by step in the meiotic recombination pathway. Adapted from Table 1 in Dapper and Payseur (2019).

| Gene | Complex | Function | Direct Interactions | Citation |
|---|---|---|---|---|
| **(A) Double strand break formation** | | | | |
| *HORMAD1* | | Associates with unsynapsed chromosomes, required for accumulation of MCD recombinosomes | *IHO1* | Fukuda et al. (2010) |
| *MEI4* | MCD Recombinosome | Component of a complex that promotes DSB formation by activating SPO11 | *REC114* | Kumar et al. (2010) |
| *REC114* | MCD Recombinosome | Component of a complex that promotes DSB formation by activating SPO11 | *MEI4, IHO1* | Kumar et al. (2018) |
| *IHO1* | MCD Recombinosome | Component of a complex that promotes DSB formation by activating SPO11 | *REC114, HORMAD1* | Stanzione et al. (2016) |
| *SPO11* | | Generates double strand breaks | | Romanienko and Camerini-Otero (2000) |
| **(B) Double strand break processing** | | | | |
| *HORMAD2* | | Associates with unsynapsed chromosomes, detects chromosome asynapsis | | |
| *MRE11* | MRN complex | Part of complex that processes newly formed DSB, trims off SPO11 | *NBS1, RAD50* | Stracker and Petrini (2011) |
| *NBS1* | MRN complex | Part of complex that processes newly formed DSB, responsible for nuclear localization of the complex | *MRE11, RAD50* | Oh et al. (2016) |
| *RAD50* | MRN complex | Part of complex that processes newly formed DSB, holds broken DNA ends together | *NBS1, MRE11* | Lamarche et al. (2010) |
| *BRCC3* | | Involved in DNA repair | | Dumont and Payseur (2011) |
| **(C) Homology search and strand invasion** | | | | |
| *DMC1* | | Mediates/catalyzes homologous chromosome pairing | *RAD51* | Tarsounas et al. (1999) |
| *RAD51* | | Mediates/catalyzes homologous chromosome pairing | *DMC1* | Cloud et al. (2012) |
| *SPATA22* | | Required for the completion of strand invasion | *MEIOB* | Xu et al. (2017) |
| *MEIOB* | | Required for the completion of strand invasion | *SPATA22* | Luo et al. (2013) |
| *MCMDC2* | | Required for the formation or stabilization of DNA strand invasion events | | Finsterbusch et al. (2016) |
| **(D) Synapsis** | | | | |
| *REC8* | Cohesion complex | Maintains sister-chromatid cohesion | | Xu et al. (2005) |
| *RAD21L* | Cohesion complex | Maintains sister-chromatid cohesion, initiates synapsis | | Lee and Hirano (2011) |
| *SYCP1* | Synaptonemal complex | Binds homologous chromosomes, transverse filament | *SYCP2* | de Vries et al. (2005) |
| *SYCP2* | Synaptonemal complex | Binds homologous chromosomes, lateral element | *SYCP1, TEX11* | Winkel et al. (2009) |
| *TEX12* | Synaptonemal complex | Binds homologous chromosomes, central element | | Hamer et al. (2006) |
| **(E) Crossover/non-crossover decision** | | | | |
| *TEX11* | | Required for the recruitment of proteins that designate crossovers | *SYCP2, SHOC1* | Yang et al. (2008) |
| *SHOC1* | | Required for the recruitment of proteins that designate crossovers | *TEX11* | Guiraldelli et al. (2018) |
| *RNF212* | | Selectively localizes to a subset of DSB, stabilizing MSH4/MSH5 | *mutS Complex* | Reynolds et al. (2013) |
| *MSH4* | mutS complex | Localizes to a subset of DSB, regulating crossover number | *MSH5* | Santucci-Darmanin et al. (2000) |
| *MSH5* | mutS complex | Localizes to a subset of DSB, regulating crossover number | *MSH4* | de Vries et al. (1999) |
| **(F) Resolution** | | | | |
| *HFM1/MER3* | | Required for correct localization of MLH1 and crossover formation | | Guiraldelli et al. (2013) |
| *CNTD1* | | Required crossover maturation and recruitment of MLH1/MLH3 | *mutL Complex* | Holloway et al. (2014) |
| *MLH1* | mutL complex | Mismatch repair gene that localizes to and resolves crossovers | *MLH3* | Baker et al. (1996) |
| *MLH3* | mutL complex | Mismatch repair gene that localizes to and resolves crossovers | *MLH1* | Lipkin et al. (2002) |

Supplementary Table 2: NCBI Reference Bird Genomes

| **Species** | **Assembly** | **RefSeq Accession** | **WGS Project Reference** |
|---|---|---|---|
| *Anas platyrhynchos* | ZJU1.0 | GCF_015476345.1 | (Li et al. 2021) |
| *Anser cygnoides domesticus* | AnsCyg_PRJNA183603_v1.0 | GCF_000971095.1 | (Lu et al. 2015) |
| *Apteryx rowi* | aptRow1 | GCF_003343035.1 | (Sackton et al. 2019) |
| *Aquila chrysaetos chrysaetos* | bAquChr1.4 | GCF_900496995.4 | (Institute 2021) |
| *Athene cunicularia* | athCun1 | GCF_003259725.1 | (Mueller et al. 2018) |
| *Calidris pugnax* | ASM143184v1 | GCF_001431845.1 | (Andersson et al. 2015) |
| *Camarhynchus parvulus* | STF_HiC | GCF_901933205.1 | (Enbody and Pettersson 2022) |
| *Catharus ustulatus* | bCatUst1.pri.v2 | GCF_009819885.2 | (Delmore et al. 2020) |
| *Columba livia* | Cliv_2.1 | GCA_000337935.2 | (Shapiro et al. 2013) |
| *Corvus moneduloides* | bCorMon1.pri | GCF_009650955.1 | (Rutz et al. 2019) |
| *Coturnix japonica* | Coturnix japonica 2.1 | GCF_001577835.2 | (Rutz et al. 2019) |
| *Cyanistes caeruleus* | cyaCae2 | GCF_002901205.1 | (Mueller et al. 2016) |
| *Dromaius novaehollandiae* | droNov1 | GCF_003342905.1 | (Sackton et al. 2019) |
| *Ficedula albicollis* | FicAlb1.5 | GCF_000247815.1 | (Ellegren et al. 2012) |
| *Gallus gallus* | bGalGal1.mat.broiler.GRCg7b | GCF_016699485.2 | (Warren et al. 2022) |
| *Hirundo rustica* | bHirRus1.pri.v3 | GCF_015227805.2 | (Secomandi et al. 2021) |
| *Lepidothrix coronata* | Lepidothrix_coronata-1.0 | GCF_001604755.1 | (Warren et al. 2016) |
| *Lonchura striata domestica* | lonStrDom2 | GCF_005870125.1 | (Mets et al. 2019) |
| *Meleagris gallopavo* | Turkey_5.1 | GCF_000146605.3 | (Dalloul et al. 2010) |
| *Melopsittacus undulatus* | bMelUnd1.mat.Z | GCF_012275295.1 | (Gedman et al. 2020) |
| *Motacilla alba alba* | Motacilla_alba_V1.0_pri | GCF_015832195.1 | (Enbody et al. 2020) |
| *Nothoprocta perdicaria* | notPer1 | GCF_003342845.1 | (Sackton et al. 2019) |
| *Numida meleagris* | NumMel1.0 | GCF_002078875.1 | (Vignal and Warren 2017) |
| *Parus major* | Parus_major1.1 | GCF_001522545.3 | (Laine et al. 2016) |
| *Phasianus colchicus* | ASM414374v1 | GCF_004143745.1 | (Wang 2019) |
| *Serinus canaria* | cibio_Scana_2019 | GCF_007115625.1 | (Gazda et al. 2019) |
| *Strigops habroptila* | bStrHab1.2.pri | GCF_004027225.2 | (Rhie et al. 2021) |
| *Struthio camelus australis* | ASM69896v1 | GCF_000698965.1 | (Zhang et al. 2014) |
| *Taeniopygia guttata* | bTaeGut1.4.pri | GCF_003957565.2 | (Rhie et al. 2021; Formenti et al. 2021) |

Supplementary Table 3: Evolutionary rates and tests for positive selection in mammals. Previously published in Dapper and Payseur (2019).

| Gene | bp | N | ω | M | M1-M2 | p-value | M7-M8 | p-value | M8a-M8 | p-value | BEB |
|---|---|---|---|---|---|---|---|---|---|---|---|
| **(A) DSB formation** | | | | | | | | | | | |
| *HORMAD1* | 1212 | 16 | 0.3036 | 7 | 0 | 1.000 | 1.795 | 0.4076 | — | — | 0 |
| *MEI4* | 1170 | 16 | 0.4332 | 7 | 0 | 1.000 | 0.005 | 0.9976 | — | — | 0 |
| *REC114* | 870 | 15 | 0.4003 | 7 | 0 | 1.000 | 5.384 | 0.0677 | — | — | 0 |
| *IHO1* | 1824 | 16 | 0.7095 | **8** | 13.061 | **0.0015** | 17.571 | **0.0002** | 14.527 | **0.0001** | **1** |
| *SPO11* | 1188 | 15 | 0.1654 | 7 | 0 | 1.000 | 4.648 | 0.0980 | — | — | 0 |
| **(B) DSB processing** | | | | | | | | | | | |
| *HORMAD2* | 981 | 15 | 0.3153 | 7 | 0 | 1.000 | 3.650 | 0.1612 | — | — | 0 |
| *MRE11* | 2136 | 16 | 0.1688 | **8** | 0.363 | 0.8342 | 11.931 | **0.0026** | 4.706 | **0.0301** | 0 |
| *NBS1* | 2289 | 15 | 0.4183 | **8** | 0 | 1.000 | 12.763 | **0.0017** | 4.087 | **0.0432** | 0 |
| *RAD50* | 3936 | 16 | 0.1006 | 7 | 0 | 1.000 | 0.301 | 0.8605 | — | — | 0 |
| *BRCC3* | 954 | 15 | 0.0602 | 7 | 0 | 1.000 | 0.250 | 0.8826 | — | —- | 0 |
| **(C) Homology search and strand invasion** | | | | | | | | | | | |
| *DMC1* | 1020 | 15 | 0.0351 | 1 | 0.488 | 0.7835 | 5.000 | 0.0821 | — | — | **1** |
| *RAD51* | 1017 | 16 | 0.0268 | 7 | 0 | 1.000 | 0 | 1.000 | — | — | 0 |
| *SPATA22* | 1101 | 16 | 0.4893 | 7 | 0 | 1.000 | 0.429 | 0.8070 | — | — | 0 |
| *MEIOB* | 1425 | 16 | 0.2341 | 7 | 0 | 1.000 | 0.665 | 0.7172 | — | — | 0 |
| *MCMDC2* | 2052 | 16 | 0.2239 | 7 | 0 | 1.000 | 0.628 | 0.7307 | — | — | 0 |
| **(D) Synapsis** | | | | | | | | | | | |
| *REC8* | 1833 | 16 | 0.3698 | **8** | 0 | 1.000 | 14.690 | **0.0006** | 5.927 | **0.0149** | 0 |
| *RAD21L* | 1686 | 15 | 0.503 | **8** | 12.124 | **0.0023** | 32.050 | **>0.0001** | 12.049 | **0.0005** | **4** |
| *SYCP1* | 3015 | 16 | 0.4337 | **8** | 8.711 | **0.0128** | 26.860 | **>0.0001** | 9.243 | **0.0024** | **3** |
| *SYCP2* | 4650 | 16 | 0.5572 | **8** | 11.584 | **0.0031** | 37.200 | **>0.0001** | 15.838 | **0.0001** | 0 |
| *TEX12* | 369 | 14 | 0.2297 | 7 | 0.0565 | 0.9721 | 1.549 | 0.4610 | — | — | 0 |
| **(E) CO/NCO decision** | | | | | | | | | | | |
| *TEX11* | 2844 | 15 | 0.8483 | **8** | 60.872 | **>0.0001** | 82.665 | **>0.0001** | 61.141 | **>0.0001** | **14** |
| *SHOC1* | 4644 | 16 | 0.6113 | **8** | 12.447 | **0.0020** | 30.561 | **>0.0001** | 15.645 | **0.0001** | 0 |
| *RNF212* | 948 | 16 | 0.5014 | **8** | 0 | 1.000 | 16.366 | **0.0003** | 5.202 | **0.0226** | **1** |
| *RNF212B* | 906 | 14 | 0.4066 | 7 | 0 | 1.000 | 0.500 | 0.7788 | — | — | 0 |
| *MSH4* | 2814 | 16 | 0.2132 | **8** | 16.608 | **0.0002** | 39.447 | **>0.0001** | 23.238 | **>0.0001** | **6** |
| *MSH5* | 2565 | 15 | 0.1642 | 7 | 0 | 1.000 | 4.214 | 0.1216 | — | — | 0 |
| **(F) Resolution** | | | | | | | | | | | |
| *MER3* | 4458 | 16 | 0.3633 | 8a | 0 | 1.000 | 12.838 | **0.0016** | 3.109 | 0.0779 | 0 |
| *CNTD1* | 1026 | 15 | 0.2496 | 7 | 0 | 1.000 | 0.936 | 0.6263 | — | — | 0 |
| *HEI10* | 831 | 15 | 0.1226 | 7 | 0 | 1.000 | 0.250 | 0.8826 | — | — | 0 |
| *MLH1* | 2313 | 15 | 0.1652 | 8a | 0 | 1.000 | 12.221 | **0.0022** | 0.280 | 0.5970 | 0 |
| *MLH3* | 4419 | 16 | 0.4444 | 7 | 0 | 1.000 | 3.757 | 0.1528 | — | — | 0 |
| *MUS81* | 1665 | 16 | 0.2124 | 7 | 0 | 1.000 | 0.628 | 0.7304 | — | — | 0 |

Supplementary Table 4: NCBI Reference Fish Genomes

| **Species** | **Assembly** | **RefSeq Accession** | **WGS Project Reference** |
|---|---|---|---|
| *Amphiprion ocellaris* | AmpOce1.0 | GCF_002776465.1 | (Tan et al. 2017) |
| *Astyanax mexicanus* | Astyanax_mexicanus-2.0 | GCF_000372685.2 | (Rohner and Warren 2019) |
| *Boleophthalmus pectinirostris* | BP.fa | GCF_000788275.1 | (You et al. 2014) |
| *Chanos chanos* | fChaCha1.1 | GCF_902362185.1 | (Wellcome Sanger Institute 2019a) |
| *Clupea harengus* | Ch_v2.0.2 | GCF_900700415.2 | (Pettersson 2019) |
| *Cynoglossus semilaevis* | Cse_v1.0 | GCF_000523025.1 | (Chen et al. 2014) |
| *Cyprinodon variegatus* | C_variegatus-1.0 | GCF_000732505.1 | (Warren and Nacci 2014) |
| *Danio rerio* | GRCz11 | GCF_000002035.6 | (Howe et al. 2013) |
| *Electrophorus electricus* | fEleEle1.pri | GCF_013358815.1 | (Myers et al. 2020b) |
| *Esox lucius* | fEsoLuc1.pri | GCF_011004845.1 | (Myers et al. 2020a) |
| *Gadus morhua* | gadMor3.0 | GCF_902167405.1 | (Wellcome Sanger Institute 2019b) |
| *Gouania willdenowi* | fGouWil2.1 | GCF_900634775.1 | (Institute” 2019) |
| *Haplochromis burtoni* | NCSU_Asbu1 | GCF_018398535.1 | (Peterson et al. 2021) |
| *Hippocampus comes* | H_comes_QL1_v1.1 | GCF_001891065.2 | (Lin 2022) |
| *Ictalurus punctatus* | IpCoco_1.2 | GCF_001660625.2 | (Liu et al. 2016) |
| *Mastacembelus armatus* | fMasArm1.2 | GCF_900324485.2 | (Wellcome Sanger Institute 2019c) |
| *Myripristis murdjan* | fMyrMur1.1 | GCF_902150065.1 | (Wellcome Sanger Institute 2019d) |
| *Oncorhynchus mykiss* | USDA_OmykA_1.1 | GCF_013265735.2 | (Gao et al. 2021) |
| *Oreochromis niloticus* | O_niloticus_UMD_NMBU | GCF_001858045.2 | (Conte et al. 2017) |
| *Oryzias melastigma* | ASM292280v2 | GCF_002922805.2 | (Kim et al. 2018) |
| *Pygocentrus nattereri* | fPygNat1.pri | GCF_015220715.1 | (Myers et al. 2020c) |
| *Salmo salar* | Ssal_v3.1 | GCF_905237065.1 | (Nome and Gillard 2022) |
| *Scleropages formosus* | fSclFor1.1 | GCF_900964775.1 | (Wellcome Sanger Institute 2019e) |
| *Takifugu rubripes* | fTakRub1.2 | GCF_901000725.2 | (Wellcome Sanger Institute 2019f) |

Supplementary Table 5: Evolutionary rates and tests for positive selection in teleost fish.

| Gene | bp | N | ω | M | M1-M2 | p-value | M7-M8 | p-value | M8a-M8 | p-value | BEB |
|---|---|---|---|---|---|---|---|---|---|---|---|
| **(A) DSB formation** | | | | | | | | | | | |
| *HORMAD1* | 1657 | 22 | 0.2188 | 7 | 0 | 1 | 1.715 | 4.243e-1 | — | — | — |
| *MEI4* | 1345 | 22 | 0.3425 | 7 | 0 | 1 | 0.573 | 7.507e-1 | — | — | — |
| *REC114* | 786 | 24 | 0.2984 | 7 | 0 | 1 | 0 | 1 | — | — | — |
| *IHO1* | 542 | 21 | 0.3602 | 7 | 0 | 1 | 1.514 | 4.690e-1 | — | — | — |
| *SPO11* | 1588 | 24 | 0.1962 | 7 | 0 | 1 | 3.453 | 1.779e-1 | — | — | — |
| **(B) DSB processing** | | | | | | | | | | | |
| *HORMAD2* | 305 | 24 | 0.1763 | 7 | 0 | 1 | 0 | 1 | — | — | — |
| *MRE11* | 700 | 24 | 0.1244 | 7 | 0 | 1 | 5.245 | 7.264e-2 | — | — | — |
| *NBS1* | 2457 | 23 | 0.2417 | 7 | 0 | 1 | 0.719 | 6.979e-1 | — | — | — |
| *RAD50* | 4123 | 24 | 0.1101 | **8** | 0 | 1 | 17.423 | **1.647e-4** | 23.606 | **>0.0001** | 0 |
| *BRCC3* | 2951 | 23 | 0.0451 | 7 | 0 | 1 | 2.656 | 0.265 | — | — | — |
| **(C) Homology search and strand invasion** | | | | | | | | | | | |
| *DMC1* | 1324 | 22 | 0.0572 | **8** | 0 | 1 | 57.485 | **>0.0001** | 4.556 | **3.281e-2** | 0 |
| *RAD51* | 1380 | 18 | 0.0272 | 8a | 0 | 1 | 10.626 | 4.928e-3 | 3.636 | 5.653e-2 | — |
| *SPATA22* | 3072 | 24 | 0.2907 | 8a | 0 | 1 | 6.203 | 4.498e-2 | 1.593 | 2.069e-1 | — |
| *MEIOB* | 2417 | 23 | 0.2088 | 7 | 0 | 1 | 0 | 1 | — | — | — |
| *MCMDC2* | 2608 | 24 | 0.2043 | 8a | 0 | 1 | 19.179 | 6.845e-5 | 1.230 | 2.674e-1 | — |
| **(D) Synapsis** | | | | | | | | | | | |
| *REC8* | 564 | 23 | 0.3072 | 7 | 0 | 1 | 1.122 | 5.707e-1 | — | — | — |
| *RAD21L* | 2209 | 23 | 0.2409 | 7 | 0 | 1 | 1.920 | 3.829e-1 | — | — | — |
| *SYCP1* | 3579 | 23 | 0.2907 | 7 | 0 | 1 | 4.397 | 1.110e-1 | — | — | — |
| *SYCP2* | 1616 | 16 | 0.2997 | 7 | 0 | 1 | 2.593 | 2.735e-1 | — | — | — |
| **(E) CO/NCO decision** | | | | | | | | | | | |
| *TEX11* | 3181 | 21 | 0.2217 | 8a | 0 | 1 | 11.604 | 3.021e-3 | 1.514 | 2.185e-1 | — |
| *SHOC1* | 305 | 14 | 0.3952 | 8a | 0 | 1 | 36.591 | 1.133e-8 | 0 | 1 | — |
| *RNF212* | 1372 | 21 | 0.2336 | 7 | 0 | 1 | 0 | 1 | — | — | — |
| *MSH4* | 2654 | 22 | 0.1130 | 7 | 0 | 1 | 0.335 | 8.457e-1 | — | — | — |
| *MSH5* | 2779 | 21 | 0.1716 | 8a | 0 | 1 | 20.329 | 3.9e-5 | 2.984 | 8.409e-2 | — |
| **(F) Resolution** | | | | | | | | | | | |
| *MER3* | 3705 | 23 | 0.1788 | 7 | 0 | 1 | 0 | 1 | — | — | — |
| *CNTD1* | 966 | 23 | 0.2901 | 7 | 0 | 1 | 1.507 | 4.706e-1 | — | — | — |
| *MLH1* | 2407 | 24 | 0.0856 | 7 | 0 | 1 | 2.681 | 2.618e-1 | — | — | — |
| *MLH3* | 3656 | 24 | 0.1865 | 7 | 0 | 1 | 0 | 1 | — | — | — |

Supplementary Table 6: Correlations between substitution rate and recombination rate, measured as XO/HCN across 7 species of birds for 29 recombination genes. Posterior probabilities are are given in parenthesis.

| | Correlation coefficient | | | Partial correlation coefficient | | |
|---|---|---|---|---|---|---|
| **Gene** | **dS - ω** | **dS - XO/HCN** | **ω - XO/HCN** | **dS - ω** | **dS - XO/HCN** | **ω -XO/HCN** |
| **(A) DSB formation** | | | | | | |
| *HORMAD1* | 0.274 (0.66) | -0.538 (0.18) | -0.492 (0.20) | 0.115 (0.58) | -0.343 (0.28) | -0.298 (0.30) |
| *MEI4* | 0.134 (0.58) | 0.466 (0.81) | 0.17 (0.61) | 0.137 (0.59) | 0.287 (0.70) | 0.107 (0.57) |
| *REC114* | -0.109 (0.42) | -0.00443 (0.51) | 0.125 (0.58) | -0.0549 (0.47) | 0.00215 (0.50) | 0.113 (0.57) |
| *IHO1* | 0.476 (0.80) | -0.072 (0.43) | 0.133 (0.58) | 0.572 (0.84) | -0.116 (0.41) | 0.193 (0.62) |
| *SPO11* | -0.312 (0.31) | 0.362 (0.81) | -0.0936 (0.42) | -0.328 (0.30) | 0.191 (0.65) | 0.0152 (0.51) |
| **(B) DSB processing** | | | | | | |
| *HORMAD2* | 0.0162 (0.50) | -0.208 (0.30) | 0.0413 (0.51) | 0.0863 (0.55) | -0.121 (0.40) | 0.0616 (0.53) |
| *MRE11* | -0.0489 (0.47) | 0.181 (0.61) | 0.016 (0.51) | -0.0132 (0.49) | 0.113 (0.58) | 0.0101 (0.50) |
| *NBS1* | 0.518 (0.82) | 0.472 (0.87) | 0.361 (0.77) | 0.527 (0.83) | 0.209 (0.65) | 0.114 (0.57) |
| *RAD50* | -0.574 (0.14) | **0.663 (0.97)** | -0.415 (0.21) | -0.458 (0.21) | 0.407 (0.80) | -0.0876 (0.44) |
| *BRCC3* | 0.0205 (0.51) | 0.39 (0.73) | 0.101 (0.55) | 0.00663 (0.50) | 0.239 (0.66) | 0.0616 (0.54) |
| **(C) Homology search and strand invasion** | | | | | | |
| *DMC1* | -0.0263 (0.48) | -0.0554 (0.46) | -0.000482 (0.50) | -0.0157 (0.48) | -0.033 (0.47) | -0.000285 (0.50) |
| *RAD51* | -0.189 (0.40) | -0.735 (0.088) | 0.148 (0.58) | -0.134 (0.42) | -0.441 (0.20) | 0.0181 (0.52) |
| *SPATA22* | 0.085 (0.54) | **0.862 (0.99)** | 0.138 (0.57) | -0.0231 (0.49) | 0.531 (0.85) | 0.125 (0.59) |
| *MEIOB* | -0.223 (0.38) | -0.103 (0.44) | 0.419 (0.73) | -0.128 (0.41) | 0.0113 (0.51) | 0.281 (0.68) |
| *MCMDC2* | 0.172 (0.61) | 0.512 (0.92) | 0.152 (0.59) | 0.146 (0.59) | 0.255 (0.72) | 0.102 (0.56) |
| **(D) Synapsis** | | | | | | |
| *REC8* | 0.937 (1) | 0.225 (0.74) | 0.235 (0.75) | 0.841 (1) | -0.0224 (0.48) | 0.0562 (0.55) |
| *RAD21L* | 0.638 (0.87) | -0.502 (0.08) | -0.388 (0.18) | 0.659 (0.89) | -0.201 (0.35) | -0.00548 (0.50) |
| *SYCP1* | -0.335 (0.29) | -0.238 (0.28) | 0.0765 (0.57) | -0.342 (0.30) | -0.115 (0.40) | 0.0109 (0.51) |
| *SYCP2* | **-0.913 (0.0029)** | -0.0758 (0.42) | 0.0344 (0.53) | **-0.93 (0.0037)** | -0.0897 (0.43) | -0.0829 (0.44) |
| TEX12 | -0.0786 (0.46) | 0.248 (0.72) | 0.0708 (0.54) | -0.0981 (0.44) | 0.128 (0.61) | 0.1 (0.56) |
| **(E) CO/NCO decision** | | | | | | |
| *TEX11* | 0.0929 (0.55) | -0.0198 (0.48) | 0.127 (0.59) | 0.124 (0.57) | -0.0212 (0.48) | 0.132 (0.59) |
| *SHOC1* | -0.051 (0.46) | -0.0826 (0.40) | 0.0572 (0.53) | -0.0318 (0.47) | -0.0378 (0.46) | 0.0678 (0.54) |
| *RNF212* | -0.0479 (0.47) | -0.00572 (0.49) | -0.0592 (0.47) | -0.00861 (0.49) | -0.00895 (0.49) | -0.0454 (0.47) |
| *MSH4* | -0.0619 (0.45) | 0.0298 (0.54) | 0.132 (0.58) | -0.0425 (0.47) | 0.0377 (0.55) | 0.13 (0.57) |
| *MSH5* | -0.0762 (0.46) | 0.0211 (0.52) | 0.0826 (0.55) | -0.0517 (0.47) | 0.0172 (0.52) | 0.0699 (0.54) |
| **(F) Resolution** | | | | | | |
| *HFM1/MER3* | 0.2 (0.64) | -0.174 (0.31) | -0.161 (0.39) | 0.207 (0.63) | -0.0617 (0.44) | -0.149 (0.41) |
| *CNTD1* | 0.366 (0.74) | 0.554 (0.87) | 0.223 (0.72) | 0.407 (0.75) | 0.481 (0.79) | -0.0719 (0.46) |
| *MLH1* | -0.00945 (0.49) | 0.0265 (0.51) | 0.0994 (0.56) | 0.0286 (0.51) | 0.0232 (0.51) | 0.0744 (0.55) |
| *MLH3* | 0.335 (0.71) | 0.23 (0.73) | 0.0704 (0.57) | 0.369 (0.73) | 0.124 (0.61) | -0.00817 (0.51) |

Supplementary Table 7: Combined evolutionary rates and model of best fit for mammals, birds, and teleost fish.

| Gene | Mammal Omega | Mammal Model | Bird Omega | Bird Model | Fish Omega | Fish Model |
|---|---|---|---|---|---|---|
| **(A) DSB formation** | | | | | | |
| *HORMAD1* | 0.3036 | 7 | 0.1896 | 7 | 0.2188 | 7 |
| *MEI4* | 0.4432 | 7 | 0.4446 | **8** | 0.3425 | 7 |
| *REC114* | 0.4003 | 7 | 0.3402 | **8** | 0.2984 | 7 |
| *IHO1* | 0.7095 | **8** | 0.4641 | **8** | 0.3602 | 7 |
| *SPO11* | 0.1654 | 7 | 0.2981 | **8** | 0.1962 | 7 |
| **(B) DSB processing** | | | | | | |
| *HORMAD2* | 0.3153 | 7 | 0.2904 | 7 | 0.1763 | 7 |
| *MRE11* | 0.1688 | **8** | 0.2445 | **8** | 0.1244 | 7 |
| *NBS1* | 0.4183 | **8** | 0.4453 | **8** | 0.2417 | 7 |
| *RAD50* | 0.1006 | 7 | 0.1724 | 8a | 0.1101 | **8** |
| *BRCC3* | 0.0602 | 7 | 0.0137 | 7 | 0.0451 | 7 |
| **(C) Homology search and strand invasion** | | | | | | |
| *DMC1* | 0.0351 | 1 | 0.034 | 7 | 0.0572 | **8** |
| *RAD51* | 0.0268 | 7 | 0.0191 | 7 | 0.0272 | 8a |
| *SPATA22* | 0.4893 | 7 | 0.5132 | **8** | 0.2907 | 8a |
| *MEIOB* | 0.2341 | 7 | 0.2094 | 8a | 0.2088 | 7 |
| *MCMDC2* | 0.2239 | 7 | 0.2066 | 7 | 0.2043 | 8a |
| **(D) Synapsis** | | | | | | |
| *REC8* | 0.3698 | **8** | 0.1391 | **8** | 0.3072 | 7 |
| *RAD21L* | 0.503 | **8** | 0.4658 | **8** | 0.2409 | 7 |
| *SYCP1* | 0.4337 | **8** | 0.4472 | **8** | 0.2907 | 7 |
| *SYCP2* | 0.5572 | **8** | 0.5487 | **8** | 0.2997 | 7 |
| *TEX12* | 0.2297 | 7 | 0.4182 | 7 | - | - |
| **(E) CO/NCO decision** | | | | | | |
| *TEX11* | 0.8483 | **8** | 0.3152 | **8** | 0.2217 | 8a |
| *SHOC1* | 0.6113 | **8** | 0.5778 | **8** | 0.3952 | 8a |
| *RNF212* | 0.5014 | **8** | 0.5242 | **8** | 0.2336 | 7 |
| *MSH4* | 0.2132 | 7 | 0.2041 | **8** | 0.113 | 7 |
| *MSH5* | 0.1642 | **8** | 0.2441 | **8** | 0.1716 | 8a |
| *HFM1/MER3* | 0.3633 | 7 | 0.3727 | **8** | 0.1788 | 7 |
| **(F) Resolution** | | | | | | |
| *CNTD1* | 0.2496 | 7 | 0.3021 | **8** | 0.2901 | 7 |
| *MLH1* | 0.1652 | 8a | 0.1754 | 8a | 0.0856 | 7 |
| *MLH3* | 0.4444 | 7 | 0.4378 | **8** | 0.1865 | 7 |

#### Supplementary Figures

Supplementary Table 8: Evolutionary rates and tests for positive selection across birds at recombination genes after removal of potential MNM sites.

| Gene | bp | N | omega | M | M1-M2 | P | M7-M8 | P | M8a-M8 | P | BEB |
|---|---|---|---|---|---|---|---|---|---|---|---|
| *MEI4* | 1724 | 29 | 0.4451 | **8** | 10.791 | **0.0045** | 14.786536 | **0.0006** | 6.999 | **0.0082** | **1** |
| *REC114* | 1776 | 29 | 0.3020 | **8** | 0 | 1.0000 | 14.17213 | **0.0008** | 4.936 | **0.0263** | **1** |
| *IHO1* | 3133 | 24 | 0.4641 | **8** | 9.767 | **0.0076** | 17.288 | **0.0002** | 10.088 | **0.0015** | **1** |
| *SPO11* | 3525 | 29 | 0.2981 | **8** | 14.256 | **0.0008** | 30.404 | **<0.0001** | 16.258 | **0.0001** | **4** |
| *MRE11* | 2103 | 28 | 0.2385 | **8** | 0 | 1.0000 | 22.494354 | **<0.0001** | 7.750 | **0.0054** | **1** |
| *NBS1* | 2533 | 29 | 0.4441 | 8a | 0 | 1.0000 | 15.193354 | **0.0005** | 3.192 | 0.0740 | 0 |
| *SPATA22* | 2398 | 29 | 0.5030 | **8** | 12.062 | **0.0024** | 21.499556 | **<0.0001** | 15.150 | **0.0001** | **3** |
| *REC8* | 2571 | 29 | 0.1391 | **8** | 17.211 | **0.0002** | 37.628 | **<0.0001** | 22.989 | **<0.0001** | **6** |
| *RAD21L* | 2906 | 29 | 0.4337 | **8** | 0 | 1.0000 | 10.594238 | **0.0050** | 4.778 | **0.0288** | 0 |
| *SYCP1* | 3577 | 26 | 0.4472 | **8** | 21.222 | **<0.0001** | 47.481 | **<0.0001** | 17.056 | **<0.0001** | **6** |
| *SYCP2* | 5508 | 26 | 0.5309 | **8** | 22.423 | **<0.0001** | 65.897896 | **<0.0001** | 31.662 | **<0.0001** | **4** |
| *TEX11* | 4555 | 29 | 0.3152 | **8** | 17.872 | **0.0001** | 41.211 | **<0.0001** | 12.249 | **0.0005** | **3** |
| *SHOC1* | 7924 | 29 | 0.5508 | **8** | 33.225 | **<0.0001** | 67.399394 | **<0.0001** | 44.382 | **<0.0001** | **12** |
| *RNF212* | 3074 | 29 | 0.5242 | **8** | 12.592 | **0.0018** | 17.743 | **0.0001** | 17.004 | **<0.0001** | **4** |
| *MSH4* | 3167 | 28 | 0.2041 | **8** | 21.487 | **<0.0001** | 57.911 | **<0.0001** | 22.879 | **<0.0001** | **4** |
| *MSH5* | 5681 | 29 | 0.1864 | 8a | 2.180 | 0.3362 | 12.329216 | **0.0021** | 3.600 | 0.0578 | 0 |
| *HFM1/MER3* | 8134 | 29 | 0.3656 | **8** | 22.639 | **<0.0001** | 47.277202 | **<0.0001** | 32.456 | **<0.0001** | **1** |
| *CNTD1* | 3168 | 29 | 0.2386 | 7 | 0 | 1.0000 | 1.779072 | 0.4108 | - | - | 0 |
| *MLH3* | 5051 | 29 | 0.4640 | **8** | 31.062 | **<0.0001** | 68.60557 | **<0.0001** | 38.745 | **<0.0001** | **11** |

Supplementary Table 9: Evolutionary rates and tests for positive selection among SHH pathway genes in mammals.

| Gene | bp | N | omega | M | M1-M2 | P | M7-M8 | P | M8a-M8 | P | BEB |
|---|---|---|---|---|---|---|---|---|---|---|---|
| *BMI1* | 3540 | 16 | 0.0532 | 7 | 0 | 1 | 4.19 | 1.233e-1 | - | - | - |
| *CCNA2* | 2748 | 16 | 0.1711 | 7 | 0 | 1 | 0.26 | 8.796e-1 | - | - | - |
| *CCNB1* | 2029 | 16 | 0.1644 | 8a | 0 | 1 | 6.55 | **3.789e-2** | 2.55 | 1.104e-1 | - |
| *CCND1* | 4238 | 16 | 0.0614 | 7 | 0 | 1 | 0.28 | 8.708e-1 | - | - | - |
| *CCND2* | 6493 | 16 | 0.0634 | 7 | 0 | 1 | 0.01 | 9.971e-1 | - | - | - |
| *DHH* | 4683 | 16 | 0.0267 | 7 | 0 | 1 | 0.47 | 7.891e-1 | - | - | - |
| *EN1* | 2427 | 16 | 0.0838 | 7 | 0 | 1 | 0.000164 | 9.999e-1 | - | - | - |
| *FOXM1* | 3507 | 16 | 0.2566 | 7 | 0 | 1 | 3.62 | 1.638e-1 | - | - | - |
| *GLI1* | 3972 | 16 | 0.1717 | 7 | 0 | 1 | 3.05 | 2.182e-1 | - | - | - |
| *GLI2* | 7136 | 16 | 1.4320 | **8** | 363.35 | **<0.0001** | 365.81 | **<0.0001** | 363.29 | **<0.0001** | **194** |
| *GLI3* | 8405 | 16 | 0.1870 | 7 | 0 | 1 | 2.11 | 3.475e-1 | - | - | - |
| *IGF2* | 5580 | 16 | 0.3142 | 7 | 0 | 1 | 0.52 | 7.731e-1 | - | - | - |
| *IHH* | 2473 | 16 | 0.0510 | 7 | 0 | 1 | 0.000194 | 1 | - | - | - |
| *MYCN* | 2613 | 16 | 0.1116 | 7 | 0 | 1 | 0.33 | 8.484e-1 | - | - | - |
| *PTCH1* | 8662 | 16 | 0.0605 | 7 | 0 | 1 | 2.06 | 3.563e-1 | - | - | - |
| *SHH* | 4650 | 16 | 0.1750 | 7 | 0 | 1 | 1.74 | 4.191e-1 | - | - | - |
| *SMO* | 3977 | 16 | 0.0539 | 8a | 0 | 1 | 8.12 | **1.723e-2** | 0.000566 | 9.810e-1 | - |
| *SUFU* | 5016 | 16 | 0.0702 | 7 | 0 | 1 | 0.0001 | 1 | - | - | - |

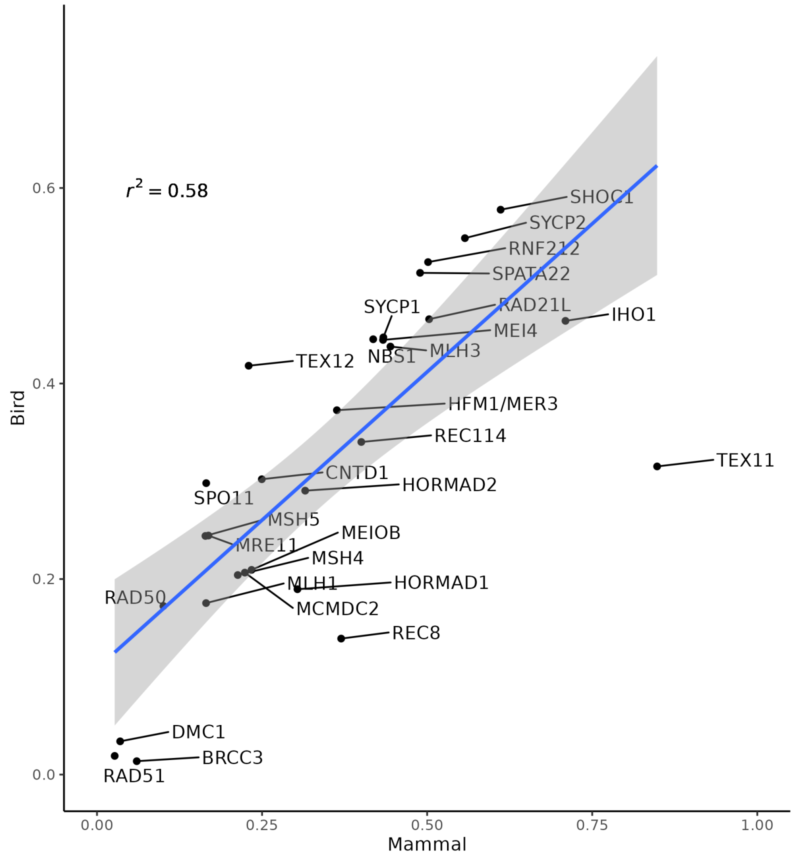


Supplementary Figure 1: Comparison of recombination omega values between birds and mammals.

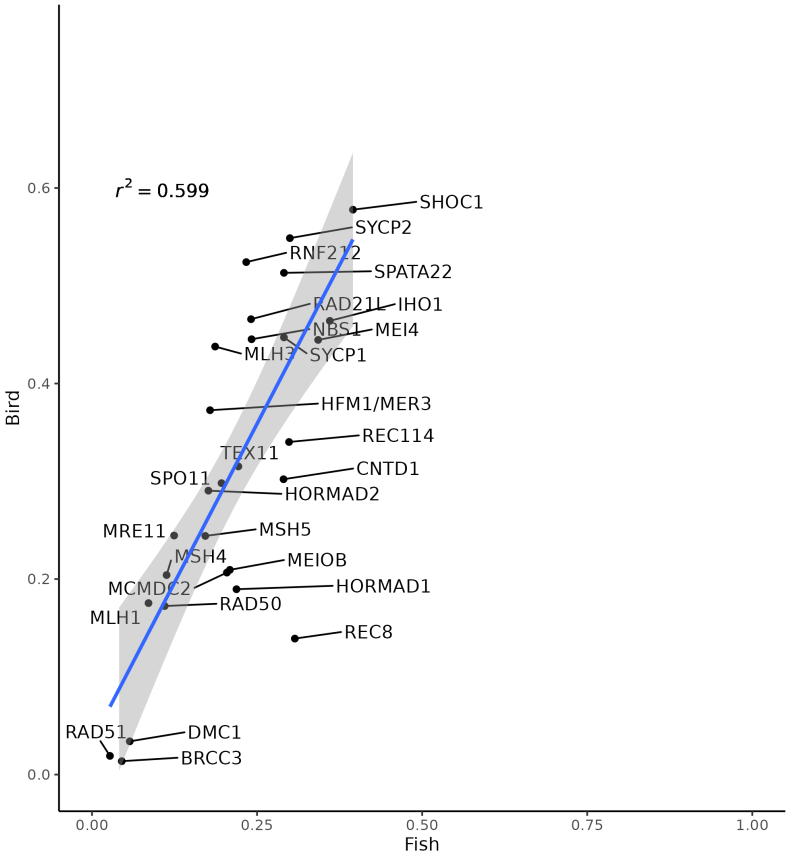


Supplementary Figure 2: Comparison of recombination omega values between birds and fish.

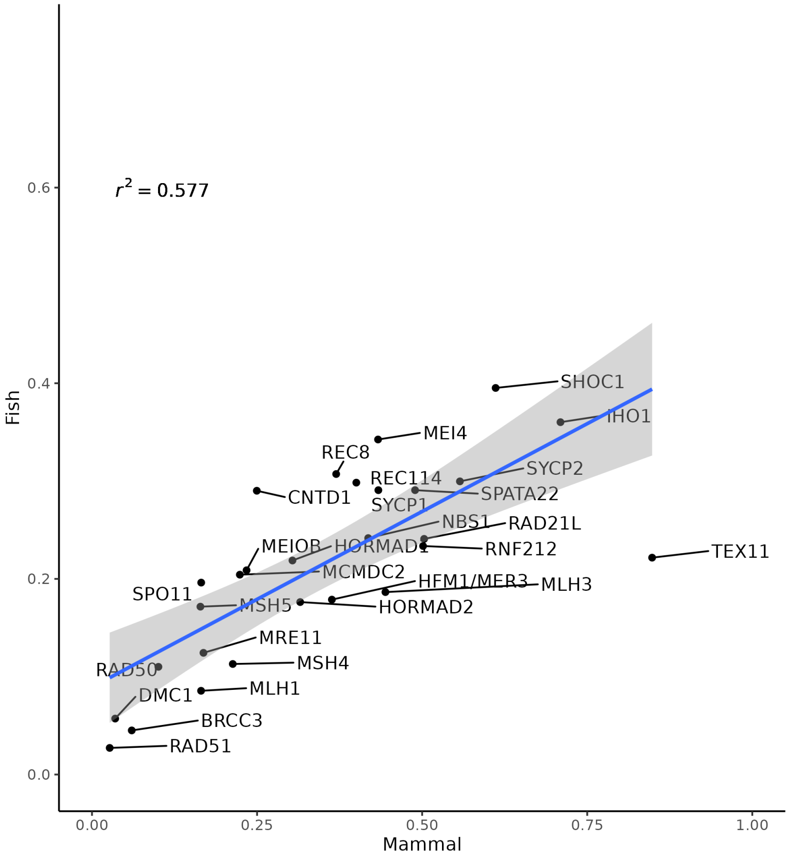


Supplementary Figure 3: Comparison of recombination omega values between mammals and fish.

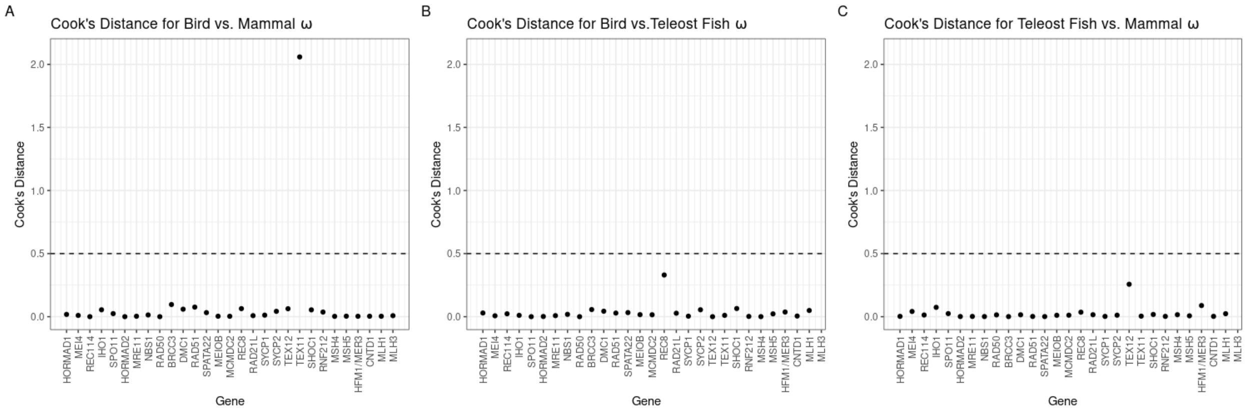


Supplementary Figure 4: Cook's Distance plots for identifying influential points in recombination gene evolutionary rate comparisons between birds and mammals (A), birds and teleost fish (B), and mammals and teleost fish (C). The significance threshold is set at 0.5 for all three comparisons.

Formenti, G., A. Rhie, J. Balacco, B. Haase, J. Mountcastle, O. Fedrigo, S. Brown, M. R. Capodiferro, F. O. Al-Ajli, R. Ambrosini, P. Houde, S. Koren, K. Oliver, M. Smith, J. Skelton, E. Betteridge, J. Dolucan, C. Corton, I. Bista, J. Torrance, A. Tracey, J. Wood, M. Uliano-Silva, K. Howe, S. McCarthy, S. Winkler,

W. Kwak, J. Korlach, A. Fungtammasan, D. Fordham, V. Costa, S. Mayes, M. Chiara, D. S. Horner, E. Myers, R. Durbin, A. Achilli, E. L. Braun, A. M. Phillippy, E. D. Jarvis, A. N. G. Kirschel, A. Digby, A. Veale, A. Bronikowski, B. Murphy, B. Robertson, C. Baker, C. Mazzoni, C. Balakrishnan, C. Lee, D. Mead, E. Teeling, E. L. Aiden, E. Todd, E. Eichler, G. J. P. Naylor, G. Zhang, J. Smith, J. Wolf, J. Touchon, K. Delmore, K. Jakobsen, L. Komoroske, M. Wilkinson, M. Genner, M. Pšenička, M. Fuxjager, M. Stratton, M. Liedvogel, N. Gemmell, P. Minias, P. O. Dunn, P. Sudmant, P. Morin, Q. Ayub, R. Kraus, S. Vernes, S. Smith, T. Lama, T. Edwards, T. Smith, T. Gilbert, T. Marques-Bonet, T. Einfeldt, B. Venkatesh, W. Johnson, W. Warren, Y. Bukhman, and The Vertebrate Genomes Project Consortium. 2021. Complete vertebrate mitogenomes reveal widespread repeats and gene duplications. Genome Biology 22(1):120. doi:10.1186/s13059-021-02336-9.

Gao, G., S. Magadan, G. C. Waldbieser, R. C. Youngblood, P. A. Wheeler, B. E. Scheffler, G. H. Thorgaard, and Y. Palti. 2021. A long reads-based de-novo assembly of the genome of the Arlee homozygous line reveals chromosomal rearrangements in rainbow trout. G3: Genes|Genomes|Genetics 11(4):jkab052. doi:10.1093/g3journal/jkab052.

Gazda, M. A., S. J. Sabatino, T. Larson, and M. Carneiro. 2019. Serinus canaria isolate sc_yellow_7_cibi, whole genome shotgun sequencing project .

Gedman, G., J. Mountcastle, B. Haase, G. Formenti, T. Wright, J. Apodaca, S. Pelan, W. Chow, A. Rhie, K. Howe, O. Fedrigo, and E. D. Jarvis. 2020. Melopsittacus undulatus isolate bMelUnd1, whole genome shotgun sequencing project .

Howe, K., M. D. Clark, C. F. Torroja, J. Torrance, C. Berthelot, M. Muffato, J. E. Collins, S. Humphray, K. McLaren, L. Matthews, S. McLaren, I. Sealy, M. Caccamo, C. Churcher, C. Scott, J. C. Barrett, R. Koch, G.-J. Rauch, S. White, W. Chow, B. Kilian, L. T. Quintais, J. A. Guerra-Assunção, Y. Zhou, Y. Gu, J. Yen, J.-H. Vogel, T. Eyre, S. Redmond, R. Banerjee, J. Chi, B. Fu, E. Langley, S. F. Maguire, G. K. Laird, D. Lloyd, E. Kenyon, S. Donaldson, H. Sehra, J. Almeida-King, J. Loveland, S. Trevanion, M. Jones, M. Quail, D. Willey, A. Hunt, J. Burton, S. Sims, K. McLay, B. Plumb, J. Davis, C. Clee, K. Oliver, R. Clark, C. Riddle, D. Elliott, G. Threadgold, G. Harden, D. Ware, S. Begum, B. Mortimore, G. Kerry, P. Heath, B. Phillimore, A. Tracey, N. Corby, M. Dunn, C. Johnson, J. Wood, S. Clark, S. Pelan, G. Griffiths, M. Smith, R. Glithero, P. Howden, N. Barker, C. Lloyd, C. Stevens, J. Harley, K. Holt, G. Panagiotidis, J. Lovell, H. Beasley, C. Henderson, D. Gordon, K. Auger, D. Wright, J. Collins, C. Raisen, L. Dyer, K. Leung, L. Robertson, K. Ambridge, D. Leongamornlert, S. McGuire, R. Gilderthorp, C. Griffiths, D. Manthravadi, S. Nichol, G. Barker, S. Whitehead, M. Kay, J. Brown, C. Murnane, E. Gray, M. Humphries, N. Sycamore, D. Barker, D. Saunders, J. Wallis, A. Babbage, S. Hammond, M. Mashreghi-Mohammadi, L. Barr, S. Martin, P. Wray, A. Ellington, N. Matthews, M. Ellwood, R. Woodmansey, G. Clark, J. D. Cooper, A. Tromans, D. Grafham, C. Skuce, R. Pandian, R. Andrews, E. Harrison, A. Kimberley, J. Garnett, N. Fosker, R. Hall, P. Garner, D. Kelly, C. Bird, S. Palmer, I. Gehring, A. Berger, C. M. Dooley, Z. Ersan-Ürün, C. Eser, H. Geiger, M. Geisler, L. Karotki, A. Kirn, J. Konantz, M. Konantz, M. Oberländer, S. Rudolph-Geiger, M. Teucke, C. Lanz, G. Raddatz, K. Osoegawa, B. Zhu, A. Rapp, S. Widaa, C. Langford, F. Yang, S. C. Schuster, N. P. Carter, J. Harrow, Z. Ning, J. Herrero, S. M. J. Searle, A. Enright, R. Geisler, R. H. A. Plasterk, C. Lee, M. Westerfield, P. J. de Jong, L. I. Zon, J. H. Postlethwait, C. Nüsslein-Volhard, T. J. P. Hubbard, H. R. Crollius, J. Rogers, and D. L. Stemple. 2013. The zebrafish reference genome sequence and its relationship to the human genome. Nature 496(7446):498–503. doi:10.1038/nature12111.

Institute”, W. S. 2019. Gouania willdenowi, whole genome shotgun sequencing project .

Institute, W. S. 2021. Aquila chrysaetos chrysaetos, whole genome shotgun sequencing project .

Kim, H.-S., B.-Y. Lee, J. Han, C.-B. Jeong, D.-S. Hwang, M.-C. Lee, H.-M. Kang, D.-H. Kim, D. Lee, J. Kim, I.-Y. Choi, and J.-S. Lee. 2018. The genome of the marine medaka Oryzias melastigma. Molecular Ecology Resources 18(3):656–665. doi:10.1111/1755-0998.12769.

Laine, V. N., T. I. Gossmann, K. M. Schachtschneider, C. J. Garroway, O. Madsen, K. J. F. Verhoeven, V. de Jager, H.-J. Megens, W. C. Warren, P. Minx, R. P. M. A. Crooijmans, P. Corcoran, B. C. Sheldon, J. Slate, K. Zeng, K. van Oers, M. E. Visser, and M. A. M. Groenen. 2016. Evolutionary signals of selection on cognition from the great tit genome and methylome. Nature Communications 7(1):10474. doi:10.1038/ncomms10474.

Li, J., J. Zhang, J. Liu, Y. Zhou, C. Cai, L. Xu, X. Dai, S. Feng, C. Guo, J. Rao, K. Wei, E. D. Jarvis, Y. Jiang, Z. Zhou, G. Zhang, and Q. Zhou. 2021. A new duck genome reveals conserved and convergently evolved chromosome architectures of birds and mammals. GigaScience 10(1):giaa142. doi: 10.1093/gigascience/giaa142.

Lin, Q. 2022. Hippocampus comes isolate QL1, whole genome shotgun sequencing project .

Liu, Z., S. Liu, J. Yao, L. Bao, J. Zhang, Y. Li, C. Jiang, L. Sun, R. Wang, Y. Zhang, T. Zhou, Q. Zeng, Q. Fu, S. Gao, N. Li, S. Koren, Y. Jiang, A. Zimin, P. Xu, A. M. Phillippy, X. Geng, L. Song, F. Sun, C. Li, X. Wang, A. Chen, Y. Jin, Z. Yuan, Y. Yang, S. Tan, E. Peatman, J. Lu, Z. Qin, R. Dunham, Z. Li, T. Sonstegard, J. Feng, R. G. Danzmann, S. Schroeder, B. Scheffler, M. V. Duke, L. Ballard, H. Kucuktas, L. Kaltenboeck, H. Liu, J. Armbruster, Y. Xie, M. L. Kirby, Y. Tian, M. E. Flanagan, W. Mu, and G. C. Waldbieser. 2016. The channel catfish genome sequence provides insights into the evolution of scale formation in teleosts. Nature Communications 7(1):11757. doi:10.1038/ncomms11757.

Lu, L., Y. Chen, Z. Wang, X. Li, W. Chen, Z. Tao, J. Shen, Y. Tian, D. Wang, G. Li, L. Chen, F. Chen, D. Fang, L. Yu, Y. Sun, Y. Ma, J. Li, and J. Wang. 2015. The goose genome sequence leads to insights into the evolution of waterfowl and susceptibility to fatty liver. Genome Biology 16(1):89. doi:10.1186/s13059-015-0652-y.

Mets, D. G., B. M. Colquitt, and M. S. Brainard. 2019. Lonchura striata domestica isolate Mets1, whole genome shotgun sequencing project .

Mueller, J. C., H. Kuhl, S. Boerno, J. L. Tella, M. Carrete, and B. Kempenaers. 2018. Evolution of genomic variation in the burrowing owl in response to recent colonization of urban areas. Proceedings of the Royal Society B: Biological Sciences 285(1878):20180206. doi:10.1098/rspb.2018.0206.

Mueller, J. C., H. Kuhl, B. Timmermann, and B. Kempenaers. 2016. Characterization of the genome and transcriptome of the blue tit Cyanistes caeruleus: Polymorphisms, sex-biased expression and selection signals. Molecular Ecology Resources 16(2):549–561. doi:10.1111/1755-0998.12450.

Myers, G., N. Karagic, A. Meyer, M. Pippel, M. Reichard, S. Winkler, A. Tracey, Y. Sims, K. Howe, A. Rhie, G. Formenti, R. Durbin, O. Fedrigo, and E. D. Jarvis. 2020a. Esox lucius isolate fEsoLuc1, whole genome shotgun sequencing project .

Myers, G., A. Meyer, O. Fedrigo, G. Formenti, A. Rhie, A. Tracey, Y. Sims, and E. D. Jarvis. 2020b. Electrophorus electricus isolate fEleEle1, whole genome shotgun sequencing project .

Myers, G., A. Meyer, N. Karagic, M. Pippel, S. Winkler, A. Tracey, J. Wood, G. Formenti, K. Howe, O. Fedrigo, and E. D. Jarvis. 2020c. Pygocentrus nattereri isolate fPygNat1, whole genome shotgun sequencing project .

Nome, T. and G. Gillard. 2022. Salmo salar, whole genome shotgun sequencing project .

Peterson, E. N., A. Elias, N. B. Roberts, and R. B. Roberts. 2021. Haplochromis burtoni strain Fernald_Lab_line, whole genome shotgun sequencing project .

Pettersson, E. M. 2019. Clupea harengus, whole genome shotgun sequencing project .

Rhie, A., S. A. McCarthy, O. Fedrigo, J. Damas, G. Formenti, S. Koren, M. Uliano-Silva, W. Chow, A. Fungtammasan, J. Kim, C. Lee, B. J. Ko, M. Chaisson, G. L. Gedman, L. J. Cantin, F. Thibaud-Nissen, L. Haggerty, I. Bista, M. Smith, B. Haase, J. Mountcastle, S. Winkler, S. Paez, J. Howard, S. C. Vernes, T. M. Lama, F. Grutzner, W. C. Warren, C. N. Balakrishnan, D. Burt, J. M. George, M. T. Biegler, D. Iorns, A. Digby, D. Eason, B. Robertson, T. Edwards, M. Wilkinson, G. Turner, A. Meyer, A. F. Kautt, P. Franchini, H. W. Detrich, H. Svardal, M. Wagner, G. J. P. Naylor, M. Pippel, M. Malinsky, M. Mooney, M. Simbirsky, B. T. Hannigan, T. Pesout, M. Houck, A. Misuraca, S. B. Kingan, R. Hall, Z. Kronenberg, I. Sović, C. Dunn, Z. Ning, A. Hastie, J. Lee, S. Selvaraj, R. E. Green, N. H. Putnam, I. Gut, J. Ghurye, E. Garrison, Y. Sims, J. Collins, S. Pelan, J. Torrance, A. Tracey, J. Wood, R. E. Dagnew, D. Guan, S. E. London, D. F. Clayton, C. V. Mello, S. R. Friedrich, P. V. Lovell, E. Osipova, F. O. Al-Ajli, S. Secomandi, H. Kim, C. Theofanopoulou, M. Hiller, Y. Zhou, R. S. Harris, K. D. Makova, P. Medvedev, J. Hoffman, P. Masterson, K. Clark, F. Martin, K. Howe, P. Flicek, B. P. Walenz, W. Kwak, H. Clawson, M. Diekhans, L. Nassar, B. Paten, R. H. S. Kraus, A. J. Crawford, M. T. P. Gilbert, G. Zhang, B. Venkatesh, R. W. Murphy, K.-P. Koepfli, B. Shapiro, W. E. Johnson, F. Di Palma, T. Marques-Bonet, E. C. Teeling, T. Warnow, J. M. Graves, O. A. Ryder, D. Haussler, S. J. O'Brien, J. Korlach, H. A. Lewin, K. Howe, E. W. Myers, R. Durbin, A. M. Phillippy, and E. D. Jarvis. 2021. Towards complete and error-free genome assemblies of all vertebrate species. Nature 592(7856):737–746. doi:10.1038/s41586-021-03451-0.

Rohner, N. and W. C. Warren. 2019. Astyanax mexicanus isolate Rio Sabinas/Rio Valles surface cross, whole genome shotgun sequencing project .

Rutz, C., C. Fungtammasan, J. Mountcastle, G. Formenti, W. Chow, K. Howe, M. P. Steele, J. Fernandes, M. T. P. Gilbert, O. Fedrigo, E. D. Jarvis, and N. Gemmell. 2019. Corvus moneduloides isolate bCorMon1, whole genome shotgun sequencing project .

Sackton, T. B., P. Grayson, A. Cloutier, Z. Hu, J. S. Liu, N. E. Wheeler, P. P. Gardner, J. A. Clarke, A. J. Baker, M. Clamp, and S. V. Edwards. 2019. Convergent regulatory evolution and loss of flight in paleognathous birds. Science 364(6435):74–78. doi:10.1126/science.aat7244.

Secomandi, S., G. Formenti, A. Rhie, O. Fedrigo, J. Mountcastle, B. Haase, J. Balacco, J. Collins, W. Chow, K. Howe, G. R. Gallo, L. Gianfranceschi, A. Bonisoli-Alquati, N. Saino, and E. D. Jarvis. 2021. Hirundo rustica isolate bHirRus1, whole genome shotgun sequencing project .

Shapiro, M. D., Z. Kronenberg, C. Li, E. T. Domyan, H. Pan, M. Campbell, H. Tan, C. D. Huff, H. Hu, A. I. Vickrey, S. C. Nielsen, S. A. Stringham, H. Hu, E. Willerslev, M. T. P. Gilbert, M. Yandell, G. Zhang, and J. Wang. 2013. Genomic diversity and evolution of the head crest in the rock pigeon. Science (New York, N.Y.) 339(6123):1063–1067. doi:10.1126/science.1230422.

Tan, M. H., H. M. Gan, Y. P. Lee, M. P. Hammer, and C. M. Austin. 2017. Amphiprion ocellaris isolate DU_AmpOce, whole genome shotgun sequencing project .

Vignal, A. and W. Warren. 2017. Numida meleagris breed g44 Domestic line isolate 19003, whole genome shotgun sequencing project .

Wang, B. 2019. Phasianus colchicus isolate SZU-A5-319, whole genome shotgun sequencing project .

Warren, W., G. Formenti, O. Fedrigo, B. Haase, J. Mountcastle, J. Balacco, A. Tracey, V. Schneider, R. Okimoto, H. Cheng, R. Hawken, K. Howe, and E. D. Jarvis. 2022. Gallus gallus breed Cross of Broiler mother + white leghorn layer father isolate bGalGal1, whole genome shotgun sequencing project .

Warren, W., B. Loiselle, M. Braun, and B. Ryder. 2016. Lepidothrix coronata voucher LSUMZ:110521 isolate B3197, whole genome shotgun sequencing project .

Warren, W. and D. Nacci. 2014. Cyprinodon variegatus isolate N-32, whole genome shotgun sequencing project .

Wellcome Sanger Institute. 2019a. Chanos chanos, whole genome shotgun sequencing project.

Wellcome Sanger Institute. 2019b. Gadus morhua, whole genome shotgun sequencing project .

Wellcome Sanger Institute. 2019c. Mastacembelus armatus, whole genome shotgun sequencing project .

Wellcome Sanger Institute. 2019d. Myripristis murdjan, whole genome shotgun sequencing project .

Wellcome Sanger Institute. 2019e. Scleropages formosus, whole genome shotgun sequencing project .

Wellcome Sanger Institute. 2019f. Takifugu rubripes, whole genome shotgun sequencing project .

You, X., C. Bian, Q. Zan, X. Xu, X. Liu, J. Chen, J. Wang, Y. Qiu, W. Li, X. Zhang, Y. Sun, S. Chen, W. Hong, Y. Li, S. Cheng, G. Fan, C. Shi, J. Liang, Y. Tom Tang, C. Yang, Z. Ruan, J. Bai, C. Peng, Q. Mu, J. Lu, M. Fan, S. Yang, Z. Huang, X. Jiang, X. Fang, G. Zhang, Y. Zhang, G. Polgar, H. Yu, J. Li, Z. Liu, G. Zhang, V. Ravi, S. L. Coon, J. Wang, H. Yang, B. Venkatesh, J. Wang, and Q. Shi. 2014. Mudskipper genomes provide insights into the terrestrial adaptation of amphibious fishes. Nature Communications 5(1):5594. doi:10.1038/ncomms6594.

Zhang, G., C. Li, Q. Li, B. Li, D. M. Larkin, C. Lee, J. F. Storz, A. Antunes, M. J. Greenwold, R. W. Meredith, A. Ödeen, J. Cui, Q. Zhou, L. Xu, H. Pan, Z. Wang, L. Jin, P. Zhang, H. Hu, W. Yang, J. Hu, J. Xiao, Z. Yang, Y. Liu, Q. Xie, H. Yu, J. Lian, P. Wen, F. Zhang, H. Li, Y. Zeng, Z. Xiong, S. Liu, L. Zhou, Z. Huang, N. An, J. Wang, Q. Zheng, Y. Xiong, G. Wang, B. Wang, J. Wang, Y. Fan, R. R. da Fonseca, A. Alfaro-Núñez, M. Schubert, L. Orlando, T. Mourier, J. T. Howard, G. Ganapathy, A. Pfenning, O. Whitney, M. V. Rivas, E. Hara, J. Smith, M. Farré, J. Narayan, G. Slavov, M. N. Romanov, R. Borges, J. P. Machado, I. Khan, M. S. Springer, J. Gatesy, F. G. Hoffmann, J. C. Opazo, O. Håstad, R. H. Sawyer, H. Kim, K.-W.

Kim, H. J. Kim, S. Cho, N. Li, Y. Huang, M. W. Bruford, X. Zhan, A. Dixon, M. F. Bertelsen, E. Derryberry, W. Warren, R. K. Wilson, S. Li, D. A. Ray, R. E. Green, S. J. O'Brien, D. Griffin, W. E. Johnson, D. Haussler, O. A. Ryder, E. Willerslev, G. R. Graves, P. Alström, J. Fjeldså, D. P. Mindell, S. V. Edwards, E. L. Braun, C. Rahbek, D. W. Burt, P. Houde, Y. Zhang, H. Yang, J. Wang, Avian Genome Consortium, E. D. Jarvis, M. T. P. Gilbert, J. Wang, C. Ye, S. Liang, Z. Yan, M. L. Zepeda, P. F. Campos, A. M. V. Velazquez, J. A. Samaniego, M. Avila-Arcos, M. D. Martin, R. Barnett, A. M. Ribeiro, C. V. Mello, P. V. Lovell, D. Almeida, E. Maldonado, J. Pereira, K. Sunagar, S. Philip, M. G. Dominguez-Bello, M. Bunce, D. Lambert, R. T. Brumfield, F. H. Sheldon, E. C. Holmes, P. P. Gardner, T. E. Steeves, P. F. Stadler, S. W. Burge, E. Lyons, J. Smith, F. McCarthy, F. Pitel, D. Rhoads, and D. P. Froman. 2014. Comparative genomics reveals insights into avian genome evolution and adaptation. Science 346(6215):1311–1320. doi:10.1126/science.1251385.
